# Decoding Proteoforms with Single Acid Resolution Using a *Sub*-nanometer Diameter Pore

**DOI:** 10.1101/2022.12.22.521660

**Authors:** Apurba Paul, Archith Rayabharam, Punam Murkute, Lisa Almonte, Eveline Rigo, Zhuxin Dong, Ashutosh Kumar, Joshy Joseph, Narayana Aluru, Gregory Timp

**Affiliations:** Electrical Engineering and Biological Science, University of Notre Dame, Notre Dame, IN 46556 USA; Department of Mechanical Engineering, University of Texas, Austin, TX 78712-1229 USA

**Keywords:** single molecule protein sequencing, nanotechnology

## Abstract

When a denatured protein isoform (i.e., a proteoform) immersed in electrolyte is impelled by an electric field through a sub-nanometer-diameter pore (i.e., a sub-nanopore) spanning a thin membrane, the sequence of amino acid (AA) residues constituting the proteoform can be directly “read” one at a time by measuring fluctuations in the electrolytic current. Corroborating this assertion, an analysis of the pore current with <u>m</u>olecular <u>d</u>ynamic (MD) simulations reveals that the fluctuations are correlated to the sequence of AA volumes, the water in the pore and affected by the acid mobility. After alignment to account for variations in the acid mobility, the simulated pore current is nearly perfectly correlated to the pattern of empirical fluctuations. To prove out the prospects for decoding proteoforms this way, site-specific <u>p</u>ost-translational <u>m</u>odifications (PTMs) and point mutations in amyloid-beta (Aβ _1-42_) are analyzed with a sub-nanopore assay. The results show that single acids can be resolved in proteoforms with a dynamic range limited by the size of phenylalanine and glycine. With this sensitivity and single acid resolution, the sequence of a scrambled variant of Aβ _1-42_ was discriminated with a p-value < 10^-5^.

---

Protein isoforms or proteoforms are proteins that are derived from closely related genes or <u>p</u>ost-translational <u>m</u>odifications (PTMs) of them. Proteoforms may have related primary structures, but they can function very differently—even antagonistically. Yet, apparently biology demands such diversity. It is estimated that there are about 100 isoforms per gene.(1–4) Since the structure dictates the function, based on these statistics, direct *de novo* sequencing of a protein seems essential to proteomics, but it is exacting work. Presently, proteomics relies mainly on <u>M</u>ass <u>S</u>pectrometry (MS)(5,6) to classify a protein and it typically requires more than a million copies to do it.(7–10) So, MS lacks the sensitivity and the dynamic range to detect less abundant peptides in a clinical specimen. Even if the peptide is correctly classified this way, the discovery of PTMs and their site assignments are still problematic; the sites are assigned statistically within a peptide.(11,12)

As an alternative to MS, it is now possible to directly “read”, one at a time, the sequence of <u>a</u>mino <u>a</u>cids (AAs) constituting a denatured proteoform immersed in electrolyte by impelling it through a *sub*-nanometer-diameter pore in a membrane a few nanometers thick, i.e. a *sub*-nanopore and measuring the pore current.(13−18) As the denatured proteoform, agglomerated with <u>s</u>odium <u>d</u>odecyl <u>s</u>ulfate (SDS), moves processively through the *sub*-nanopore, the SDS is stripped from the proteoform(14) and the ion flow around it fluctuates reproducibly in relation to the acids in the pore waist. If the pore current is sampled at a high enough rate (Ms/sec) in line with the acid velocity, and the electric field in the pore is confined to a *sub*-nanometer extent near the waist, then the fluctuations in the pore current can inform on a single acid and the pattern of fluctuations reflects the AA sequence.

Several innovations have come together to put this alternative into practice. First, because the average volume of an AA residue is only about 0.15 nm^3^, the amplitude of a fluctuation in the blockade scales like the molecular volume in the waist. So, a pore with *sub*-nanometer dimensions is indicated to recover a signal. Moreover, to recover the signal from a single acid, the electric field and therefore the current density must be tightly constrained to *sub*-nanometer dimensions consistent with the distance between AAs, which translates to a *sub*-nanopore in a nanometer-thick membrane. Second, when the pore current is sampled at a high rate, the signal is noisy due to the wide bandwidth. So, to reduce the noise, the signal has to be filtered and averaged over clusters containing hundreds of blockades with similar fluctuation patterns.(19) The fluctuation patterns are identified using an <u>a</u>ffinity <u>p</u>ropagation (AP) algorithm without reference to any model for the peptide structure, unlike earlier work that compared the fluctuation patterns to volume models of the sequence.(13−16) This distinction is important because, beside the Nyquist criterion for sampling the electrical signal, there are no assumptions about fluctuation amplitude and timing, and it opens a window into the identification of a protein in a mixture. Third, to call acids from the fluctuation patterns in the pore current, <u>m</u>olecular <u>d</u>ynamic (MD) simulations that are accomplished with atomic precision are indispensable. An analysis of the pore current with MD reveals that the fluctuations are correlated to the sequence of AA volumes, the water in the pore and affected by the acid mobility. Thus, MD can be used as basis for interpreting the fluctuation pattern with confidence, which is unprecedented. Finally, after accounting for differences in the acid mobility by using <u>d</u>ynamical time <u>w</u>arping (DTW) to align the time sequence,(16, 20−22) the empirical fluctuations in the pore current are nearly perfectly correlated to the MD simulations of the ion flow around a single acid translocating through the pore. The correlation between the aligned time sequence and the MD simulations of the current means that the acid sequence can be “read” with single acid resolution from the fluctuations with only a minimal amount of material—a few hundred molecules—required to identify a peptide. “Reading” a peptide with this resolution from so few copies holds out the prospect of transforming proteomics.

To prove out the prospects for calling acids this way, amyloid-beta peptide (Aβ_1-42_) along with variants of it were assayed with a *sub*-nanopore. Aβ_1-42_ is the predominant species of <u>a</u>myloid-<u>β</u> (Aβ) affecting the pathology found in the brain of people with <u>A</u>lzheimer’s <u>d</u>isease (AD) and Down’s syndrome. Aβ is supposed to accumulate in AD due to an imbalance in the production and clearance of the peptide resulting in the formation of amyloid plaques. Since there is no apparent change in the production of the Aβ peptide in AD, the propensity to form toxic species is likely driven by other factors besides just the primary structure of Aβ, such as the terminal acids, mutations and PTMs.(23−30) Importantly, aromatic residues like tryptophan (W) and phenylalanine (F) seem to drive amyloid formation *in silico*, but even non-aromatic acids such as cysteine (C) and methionine (M) reportedly produce amyloid-like structures.(28,29) Moreover, several PTMs have also been implicated in an increase in the aggregation rate— among them are phosphorylation of S8 and S26(24) and glycosylation.(30)

## *Sub*-nanopore fabrication and characterization

Following other work (13–18), a *sub*-nanopore was sputtered through the *a*-*Si* only a few micrometers on-edge with a thickness that ranged from *t* = 3.5 to 6.0 nm (Figs. 1a,b) using a tightly focused, high energy (300 keV) electron beam formed in a <u>s</u>canning transmission <u>e</u>lectron <u>m</u>icroscope (STEM, Methods). Sometimes immediately after sputtering, the pore was visualized *in situ* with TEM to reveal a 1.5 nm-diameter at the waist defined by the shot noise (Fig. 1c). However, when the same pore was re-acquired and the topography visualized with <u>H</u>igh-<u>A</u>ngle <u>A</u>nnular <u>D</u>ark <u>F</u>ield (HAADF) using an aberration-corrected STEM after exposure to the ambient for 1−3 da, a smaller pore diameter was observed (Fig. 1d). According to the line profile, the pore topography had a steep cone-angle > 5-15° that broadened to >16−30° near each orifice with an irregular waist > 0.40 nm in diameter generally. The pore topography was bi-conical, but since the STEM images revealed only the mass density under the probe beam, the angles indicated in the line profile represented twice the actual cone-angles.(18) According to this evidence then, the pore topography was likely affected, not only by electron-beam sputtering, but also by oxidation of silicon in the ambient.(31–34) However, even though the STEM images were acquired at low voltage (80 kV) and/or with low beam current (16 pA), it was still possible that sputtering induced by the probe beam backfilled the pore.

**FIGURE 1.**
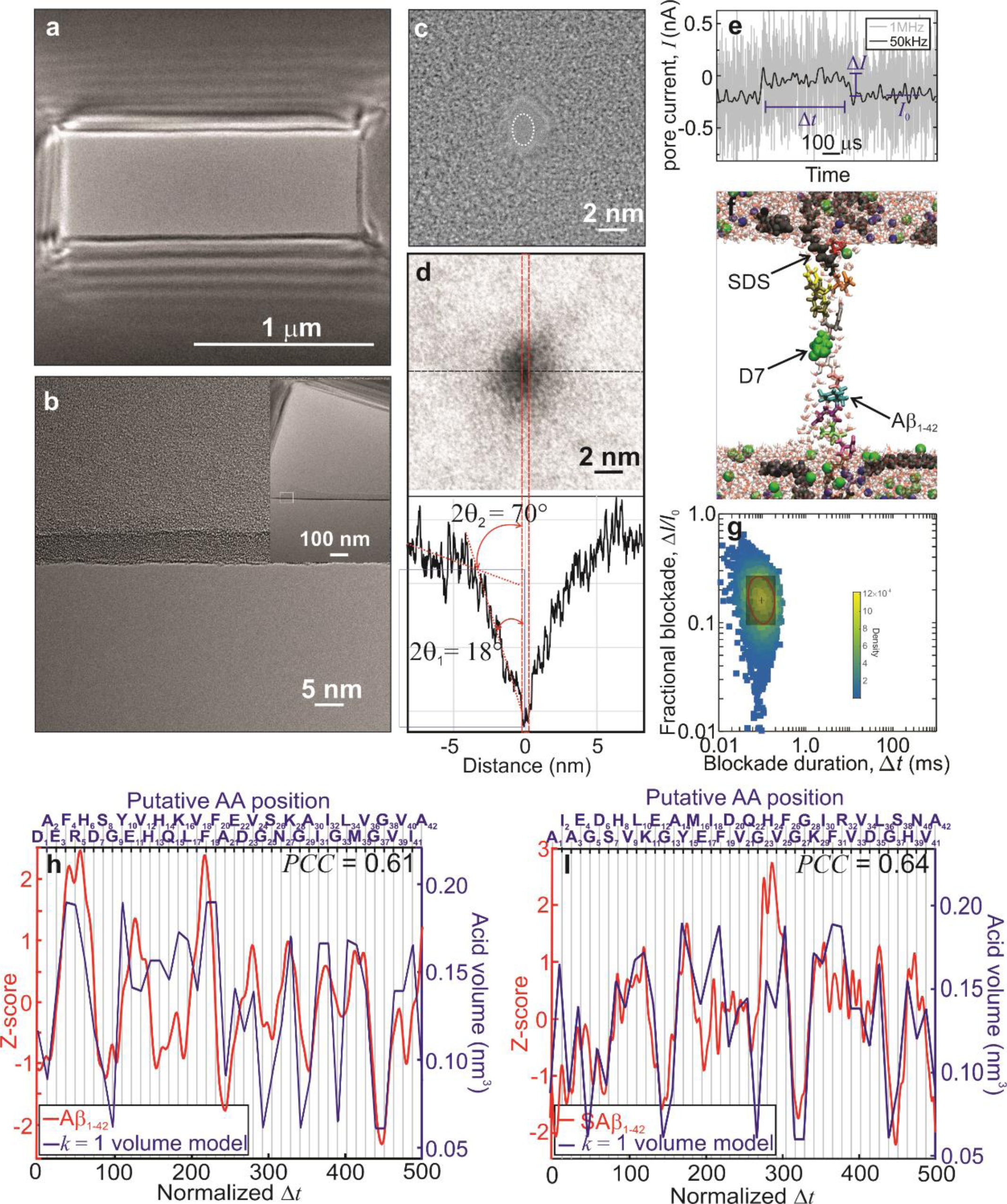
“Read” resolution and fidelity using a *sub*-nanopore through a thin *a*-*Si* membrane. **(a)** A transmission <u>e</u>lectron <u>m</u>icrograph (TEM) is shown of a thin *a*-*Si* membrane spanning a 0.56 μm × 1.59 μm window through a 100 μm thick silicon handle wafer. **(b)** A TEM of an edge in a thin *a-Si* membrane after it was purposefully punctured to reveal a 5.3 nm thickness. The *a*-*Si* membrane was nominally 5 nm thick, but typically the thickness ranged from 3.5 to 6 nm. **(b, Inset)** A low magnification micrograph is shown of the fold in the membrane (b). **(c)** A TEM image taken *in situ* of a pore immediately after sputtering through a *a*-*Si* membrane. The cross-section of the pristine pore was estimated from the shot noise associated with electron transmission through the pore to be about 1.5 nm × 2.1 nm (dotted circle). **(d, top)** A HAADF-STEM image, acquired with an aberration-corrected STEM, is shown of the pore in (c) after exposure to the ambient. **(d, bottom)** The profile was taken along the dashed (horizontal) line in (d, top). The pore cross-section shrunk to about 0.4 nm × 0.7 nm likely due to the growth of a native oxide. The angles *θ* _1_ and *θ* _2_ reflect twice the cone angle of the bi-conical topography. **(e)** A current trace is shown that illustrates the blockade current associated with translocations of single molecules of Aβ_1-42_ through a *sub*-nanopore measured with an applied voltage of 0.6 V. The pore current was amplified over a bandwidth of 1.8 MHz, filtered with a low pass 1 MHz filter and sampled at 3.6 MHz (gray line). Another version of the same data, smoothed with a low-pass 50 kHz eight-pole Bessel filter (black line), is juxtaposed on the 1 MHz data. The definition of the blockade current, Δ*I*, the blockade duration, Δ*t*, and the open pore current, *I*_0_, are indicated. According to these definitions, higher current values correspond to larger blockade currents. **(f)** A <u>v</u>isual <u>m</u>olecular <u>d</u>ynamics (VMD) snapshot is shown depicting the translocation of Aβ_1-42_–SDS aggregate forced through a bi-conical *sub*-nanopore with a cone angle of 10° near the waist, spanning a 4.0 nm thick *SiO*_2_ membrane. The diameter at the waist was 0.4 nm. The pore is ghosted in the figure; only the peptide, the SDS, the *Na*^+^ (yellow/green) and *Cl*^−^ (cyan) ions and the water molecules are represented. Due to the steric hindrance, the SDS is stripped from the peptide near the pore waist. **(g)** A heat map is shown that illustrates the distribution of blockades relative to the open pore current (Δ*I/I*_0_) versus the blockade duration (Δ*t*) associated with single denatured Aβ_1-42_ molecules translocating through a *sub*-nanopore acquired at 0.6 V. The red contour line captures 50% of the population and the red cross-hairs depict the position of the median. The region highlighted in red indicates that portion of the distribution used to form the consensus. **(h)** A plot is shown of a blockade current consensus of 806 blockades versus normalized duration measured when denatured Aβ_1-42_ was electrically forced through a *sub*-nanopore (red line). The signals forming the consensus were acquired at 0.6 V amplified over a 1.8 MHz bandwidth, sampled at 3.6 MS/sec and digitally filtered to reveal the fluctuations. Juxtaposed with the empirical data is the corresponding volume model for the peptide (assuming *k* = 1, blue line). The blockade current was well correlated (*PCC* = 0.61) to the volume model. **(i)** Like (h), a plot is shown of a blockade consensus of 975 blockades versus normalized duration acquired by electrically forcing a scrambled variant of Aβ_1-42_, SAβ_1-42_, through the same *sub*-nanopore as in (h) (red line). Juxtaposed with the empirical data is the corresponding volume model (assuming *k* = 1, blue line). This empirical consensus was also well correlated (*PCC* = 0.64) with the volume model.

<u>F</u>inite <u>e</u>lement <u>s</u>imulations (FES) that mimic the pore topography while accounting for the finite-size of the ion, the collapse of the electrolyte mobility and the negative surface charge were used to estimate the electric field distribution and electrolytic conductance through a *sub*-nanopore immersed in electrolyte.(17) FESs revealed an electric field focussed to *sub*-nanometer dimensions near the waist (supplemental Figs. 1a,b). Depending on the cone-angle and membrane thickness, the electric field was supposedly focussed into a region about 0.70 nm in extent, which is about the size of a single stretched acid residue (0.5−0.55 nm).(14) The correspondence between the empirical conductance and the FESs lent credence to this idea (supplemental Fig. 1c). So, if only one (or two) acid(s) occupied the waist at a time, it was reasoned that the blockade current would mainly measure that residue.

These assertions were corroborated by direct measurements of the force and the pore current as a single Aβ_1-42_ molecule, decorated with SDS and tethered to the tip of an <u>a</u>tomic force <u>m</u>icroscope (AFM) cantilever, was impelled through a *sub*-nanopore spanning a nominally 5 nm thick *a-Si* membrane (supplemental Fig. 2). SDS is an anionic detergent that works, in combination with heat (45−100°C) and reducing agents like <u>β</u>-<u>m</u>ercapto<u>e</u>thanol (β−ME), to denature proteins. The SDS molecule end-to-end is about 1.7 nm long and 0.25 nm wide. Ostensibly, it binds regularly along the length of the protein backbone and imparts a nearly uniform negative charge that swamps the intrinsic charges in the peptide, disrupting the native folded structure and producing a linear “rod-like” aggregate.(35) This is important because the acids constituting a peptide are not uniformly charged and so, the uniform charge on the protein-SDS aggregate is supposed to afford some control over the translocation kinetics through the electric force that develops when a voltage is applied across the membrane (36,37). The electric field then impels the aggregate progressively, “frictionlessly” through the pore with a nearly uniform velocity, which facilitates “reading” the acid sequence. However, according to MD simulations, the SDS typically penetrates only about 1.2−1.3 nm into the pore during the translocation of the peptide across the membrane. So, due to the size of the aggregate, which is about 1.2−1.4 nm wide, the SDS is cleaved off by the steric constraints imposed by the pore topography leaving just a few acids exposed in the waist. These MD simulations are consistent with force measurements with and without SDS on the *trans*-side of the membrane performed with AFM (Fig. 1f).(14) That data revealed that the blockade was relieved (on average) about 1 nm before the electric force on the molecule terminated. This observation was interpreted as evidence that the current blockade occurs within about 1 nm of the waist since the electric field hardly extends beyond that (supplemental Fig. 2). Importantly, the translocation kinetics are not always “frictionless”. AFM measurements have revealed that the peptide can also progress through the *sub*-nanopore in slip-and-stick motion introducing delays in the timing that frustrate “reading” the sequence.(14)

Finally, it has been shown empirically and corroborated with MD that the ion mobility collapses as the diameter of the *sub*-nanopore shrinks(17) and so, it was reasoned that it would also affect the acid mobility and velocity in the same way.(36) Doubtless other AAs outside the waist still contributed at least marginally to the blockade current and the acid mobility in the pore, however.

## Detecting acid residues in measurements of the blockade current

When a nearly pH-neutral (pH 6.6 ± 0.1) solution containing the denatured peptide was introduced on the *cis*-side of a *sub*-nanopore with a voltage of 0.4−0.7 V applied across the membrane, blockades (defined as: Δ*I* ≡ *I* – *I*_0_) were observed in the open pore current, *I*_0_ (Fig. 1e). Some, but not all, of these blockades were attributed to the translocation of a single peptide molecule across the membrane through the *sub*-nanopore (Fig. 1f). To interrogate each acid in the peptide, the pore current, *I*, was amplified over a wide bandwidth and sampled at a high rate (Methods). However, the resulting signal was partially obscured by random electrical noise (Fig. 1e), and so, to identify the blockades, the current traces were passed through a low-pass digital filter. To facilitate the analysis, the blockades were classified by the fractional change in the pore current relative to the open pore value (Δ*I/I*_0_) and the duration of the blockade (Δ*t*). The aggregate data were then represented by a normalized heat map of the <u>p</u>robability <u>d</u>ensity function (*PDF*) reflecting the density and distribution of blockades (Fig. 1g). For example, the median blockade for the distribution delineated by the crosshairs in Fig. 1g was at: Δ*I/I*_0_ = 0.16, Δ*t* = 100 μsec; the red contour captured 50% of the population.

Regardless of the length of the peptide chain, based on the median blockade duration, a peptide progresses through a *sub*-nanopore at an average rate of about one acid in 2 μsec. As an initial assessment of the “read” accuracy of a single blockade at this rate, current blockades sampled at 250 kS/sec and amplified over a bandwidth of 50−100 kHz, associated with the translocation of three block copolymers: *block-p*(R)_10_-*co-*(G_3_S)_3_-G, 23 AAs long (denoted by R-G-S); *block-p*(K)_10_-*co-*(G_3_S)_3_-G, 23 AAs long (denoted by K-G-S); and *block-p*(K)_24_-*co-*(A_8_P_8_)_2_-A_8_, 48 AAs long (denoted by K-A-P) were tested. These three block copolymers were used as positive controls because it was reasoned that the step in acid volume would show up as an abrupt change in the pore current (supplemental Figs. 3a-c). Scrutiny of select, single blockades from these controls revealed fluctuations consistent with the variations in the acid volume between *block-p*(R) and *co-*(G_3_S)_3_-G, and *block-p*(K) and *co-*(G_3_S)_3_-G and *co-*(A_8_P_8_)_2_ in the copolymer, but despite the narrow bandwidth, the signal was still noisy and the resolution blurred. Moreover, not all the blockades examined produced precisely the same fluctuation patterns; some likely did not correspond to translocations at all, but rather the peptide straddling the pore. So, further improvement in sorting and signal averaging was indicated.

Subsequently, measurements of the fluctuations in the blockade current through a *sub*-nanopore were used to analyze eight different peptides (in addition to the three block co-polymers used for controls): a 42-residue (human) amyloid-β (Aβ_1-42_)−protein fragment and seven variants of it (supplemental Table 2). The blockade current distribution was important for accurate signal reconstruction because: the fractional change in current has been linked to the ratio of the molecular volume to the pore volume (13,38); the median duration of a blockade was a measure of the average velocity through the pore; the width of the blockade duration distribution relative to the median was a gauge of Brownian diffusion and backward fluctuations or slip-and-stick acid motion through the pore.(39)

To improve the signal-to-noise ratio, a subset of the empirical data was selected for signal averaging by culling blockades with a duration too short to satisfy the Nyquist criterion, which dictates that each acid be sampled twice, or too long to preclude back-stepping and slip-and-stick motion (Methods). Likewise, blockades with a fractional blockade current too high (Δ*I/I*_0_ > 0.3) or too low (Δ*I/I*_0_ < 0.10) to be identified with a single molecule occluding the pore were also culled (Fig. 1g, box highlighted in red).(Methods) The mean of each of the blockades in the subset was then zeroed and the result normalized by the standard deviation (*σ* ) to form the Z-score. The Z-score is an important statistic because it relates the number of standard deviations away from the mean blockade current, where the standard deviation measures the root <u>m</u>ean <u>s</u>quare (rms) of the current noise and signal together. Finally, to recover reproducible fluctuations from random noise, two common signal-averaging techniques were employed: low-pass digital filtering; and signal averaging over clusters of blockades with similar fluctuation patterns identified with an AP algorithm.(Methods) It was observed that the rms-current noise in an open pore, which is equivalent to *σ* when the mean current is suppressed, averaged over durations comparable to a blockade, diminished as 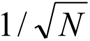 where *N* denotes the number of events with a duration indicative of random noise (supplemental Fig. 3e). However, unlike random noiser, the fluctuations in the blockade current persisted even after signal averaging with an amplitude swinging between 5−6*σ* (Methods, supplemental note #2 and supplemental Figs. 3d-f).

Using the same *sub*-nanopore for the assay, differences in the fluctuation patterns between Aβ_1-42_ and a scrambled variant of it (denoted as SAβ_1-42_), consisting of a different sequence of the same residues (supplemental Table 2), were conspicuous.(16) On comparison, it was discovered that the empirical data were correlated with naïve volume models of the two peptides (Figs. 1h,i; blue lines) derived by translating the sequence of AA volumes ascribed to the primary structure obtained from crystallography data (supplemental Table 1),(40) assuming that the blockade current corresponded to convolution over the sequence of volumes with a window *k =* 1−acid wide moving with a uniform velocity through the pore.(13) Strikingly, the amplitude of the fluctuations tracked closely the *k* = 1 naïve volume models for Aβ_1-42_ and the scrambled variant of it as indicated by the <u>P</u>earson <u>C</u>orrelation <u>C</u>oefficients (*PCCs*) *PCC* = 0.61 and 0.64, respectively. (These *PCCs* were extraordinary; typically, *PCC* = 0.4−0.5.) Yet, the empirical data for SAβ_1-42_ were poorly correlated to the volume model for Aβ_1-42_ (*PCC* = − 0.050) and the Aβ_1-42_ empirical consensus (*PCC* = 0.034). So, even though the number and constituency of AAs in the peptide did not change, the pattern of fluctuations did, and the pattern was correlated to volume models proving that the amplitudes were a measure of the acid sequence.

These results were especially relevant to the prospects for *de novo* sequencing of a protein for three reasons. First, an AP algorithm was used to identify blockades with similar fluctuation patterns regardless of the fluctuation timing or amplitude.(16) Second, the differences evident between the consensuses representing Aβ_1-42_ and SAβ_1-42_ indicate that the fluctuations patterns informed on the AA sequence of the respective peptides, which was consistent with earlier reports.(13−16) This meant that the entire molecule could be sequenced at once this way, in principle, without resorting to enzymatic digestion. Third, the correspondence between empirical consensus and the volume models proves that the fluctuations can be interpreted as high fidelity “reads” with (nearly) single residue resolution. However, it is unlikely that all twenty AAs could be discriminated unambiguously based only on a volume model alone. (For example, leucine and isoleucine have practically identical volumes.) So, the correlation between the empirical consensuses and the corresponding volume models used for calling acids was still imperfect. Importantly, the underlying assumption in the comparison of the empirical data with the volume models was that the peptides moved through the pore with a uniform velocity, which seemed suspect because it discounted the possibility of variations in the acid mobility in the pore.

## MD simulations of the translocation of Aβ_1-42_

To unravel the various components that contributed to the amplitude and timing of the fluctuations such as the acid volume, charge, mobility and the effect of the neighboring acids and nearby water molecules, the pore current associated with the translocation of Aβ_1-42_ and variants of it were simulated with MD using a model of the *sub*-nanopore topography (Fig. 2a and supplemental VideoAB1-42e) (Methods). These predictors were mutually related. For example, the fluctuation amplitude depends on the AA, which was affected by the acid volume relative to the size of the pore, the charge and the interactions between the molecule and the pore surface, coupled with the dynamics of the counter-ions, the neighboring acids, and water there.(41) To economize on computing time, peptide fragments one to four acids long occupied the pore waist at a time. This made sense because, due to the size of the SDS-peptide aggregate, the SDS must be cleaved off in the pore waist exposing just a few acids.

**FIGURE 2.**
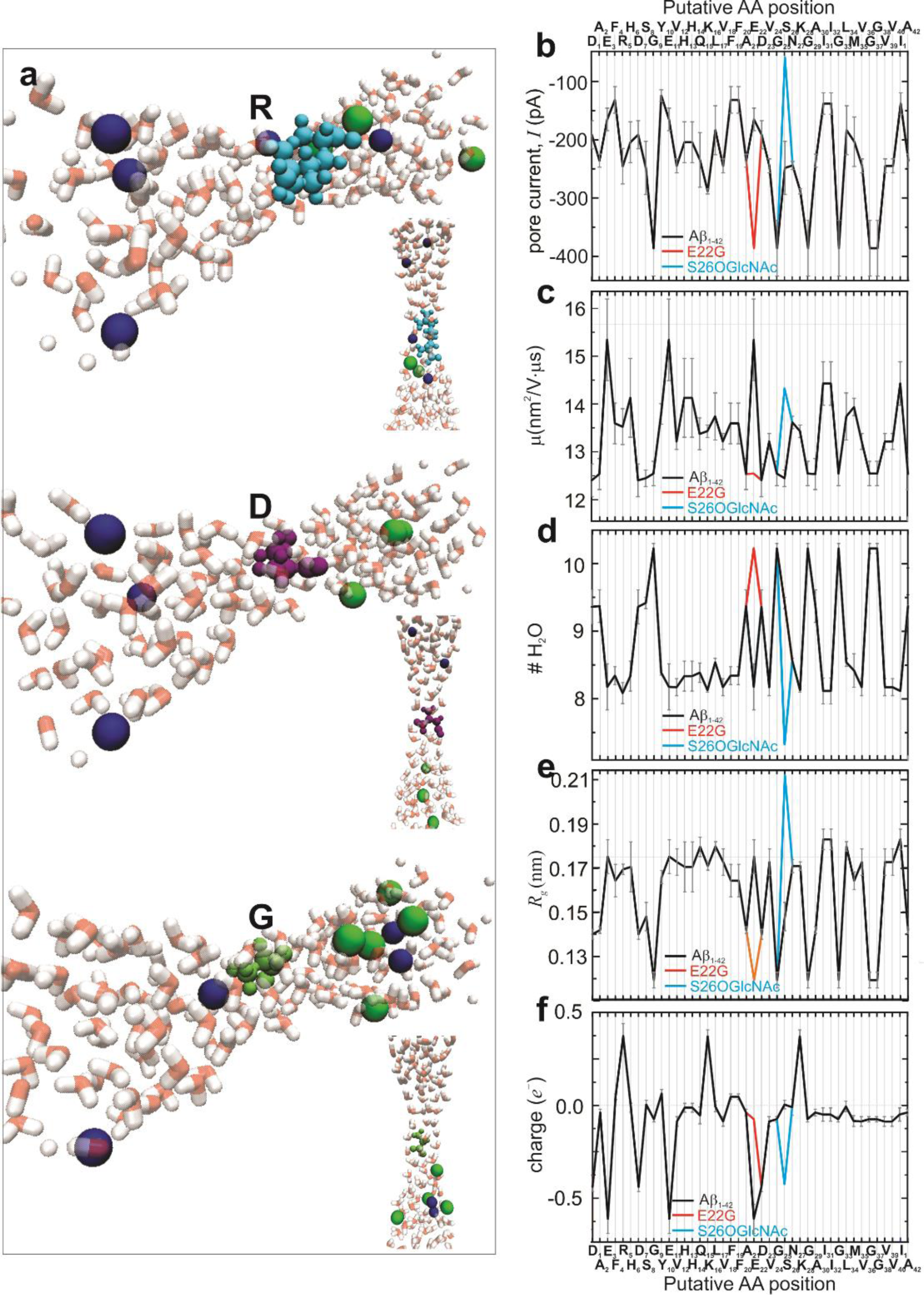
MD simulations of the translocation of Aβ_1-42_ through a *sub*-nanopore in silica. **(a)** <u>V</u>isual <u>m</u>olecular <u>d</u>ynamics (VMD) snapshots are shown depicting three amino acid residues, arginine (R), aspartic acid (D) and glycine (G) translocating through a *sub*-nanopore. The *sub*- nanopore that spanned a 4.0 nm thick membrane was defined by a bi-conical topography with a cone angle of θ_1_ = 10° and a 0.4 nm diameter at the waist. The pore is ghosted in the figure; only the water molecules, the peptide fragments and the *Na*^+^ (yellow/green) and *Cl*^−^ (blue) ions are represented. **(b)** The pore current derived from the MD simulations of the acids comprising the Aβ_1-42_ (black) peptide impelled through the *sub*-nanopore are shown for each residue. **(c,d,e,f, black lines)** Like (b), the mobility, the number of waters in the neighborhood of the acid, the gyradius (*R_g_*) and the charge extracted from MD simulations are shown, respectively.

The translocation through the pore was governed by several factors: the imposed (zero-acceleration) molecular velocity (5×10^-6^ *nm/ps*); the 5 V difference across the membrane; and the non-bonding interactions of the peptide with the pore. The (5 V) voltage drop across the membrane was important for three reasons: **1.** it more closely mimicked the conditions of the experiments; **2.** the electric force due to voltages > 5 V distorted the acid volume in the pore, which destroyed the correlation to the empirical data; and **3.** too few ions flowed around an acid residue in the pore waist for simulations performed at a lower voltage (< 0.5 V), which compromised the statistical evaluation of the current. Although the course of these simulations was similar to the AFM experiments in which a single Aβ_1-42_ molecule was impelled through a *sub*-nanopore (supplemental Fig. 2), the duration of the empirical blockade (lasting ∼30 sec at an applied voltage of 0.7 V), the measured force (∼25 *pN*), and the molecular velocity were not captured precisely in MD.

Just like the empirical data, the properties of Aβ_1-42_ calculated this way all fluctuated in correspondence with the acids in the pore waist (Figs. 2b-f, supplemental Table 1). The (negative of the) pore current was correlated to the AA volume (*PCC* = 0.77) and the gyradius, *R*_g_, (*PCC* = 0.72) that was calculated from the positions of van der Waals models for the atoms in the waist, the acid mobility (*PCC* = 0.60), and the number of waters near the acid (*PCC* = 0.71), but hardly related to the acid charge (*PCC* = 0.20). Likewise, the correlation between the simulated pore current and the acid volume extended to the variants. Coincidently, the magnitude of the pore current increased with the substitution of G for E22 (denoted by E22G) and decreased with the glycosylation of S26 (denoted by S26OGlcNAc), in correspondence with the changes in volume, mobility and waters in the pore. Specifically, corresponding to an increase (decrease) in the number of the waters in the pore waist, the mobility was diminished (enhanced) and the magnitude of the pore current swelled (shrunk).

Generally, acid residues surrounded by a larger number of water molecules (such as A, S, D and G) tended to have lower mobility, whereas acids surrounded by less water (K, R, F and I) showed higher mobility (assuming a linear fit, R^2^ = 0.54). Also, acids with a larger volume (like W compared to G) squeezed out the water in the pore (assuming a linear fit to *R_g_*, R^2^ = 0.92). The correlation between the mobility, the waters near the acid and the electrolytic pore current was rationalized by considering all the forces on the ions in the miniscule pore volume. As the peptide was impelled through the pore waist by the electric force, it dragged ions with it. Additional water in the neighborhood created drag that, in turn, slowed the residue. This coordination only happened because the ions, the water and the peptide (fragments) were constrained in the pore.

Importantly, the simulated pore current was also correlated to the empirical blockade current (*PCC* = 0.41). However, in this naïve analysis, the correspondence with the empirical data suffered because it was assumed that the velocity of the peptide through the pore was uniform; it did not account for the variations in the acid mobility. The acid mobility ranged from 12.4×10^-3^ to 15.3×10^-3^ *nm*^2^/*V·ns*, depending on the acid. So, if the speed of the acid through the pore was boosted or slackened, then the position in the time trace, *τ_i_*, and duration of the *i*^th^ fluctuation in the blockade current, Δ*τ_i_ = τ_i_ − τ_i−_*_1_, would be affected by the mobility. To appreciate the disparities, Δ*τ_i_*, was calculated for S26 (*μ*_S26_ = 12.5×10^-3^ *nm*^2^*/V·ns*) and E22 (*μ*_E22_ = 15.3×10^-3^ *nm*^2^*/V·ns*) subjected to the same electric field and then compared. Accordingly, whereas the average duration of a fluctuation per acid would be about Δ*τ* = 2.4 *μs*, Δ*τ _S_*_26_ = 2.7 *μs* for S26 and Δ*τ _E_* _22_ = 1.3 *μs* for E22. Thus, the maximum deviation in the delay Δ*τ_i_*, was about half the average.

Two conclusions follow from these simulations. First, the pore current was correlated to the size of the acid, the waters surrounding the residues in the pore waist and affected by the mobility. Generally, the magnitude of the pore current decreased (increased) as the mobility increased (decreased), as the number of water molecules decreased (increased), and as the radius of gyration increased (decreased). The variation in acid mobility would undoubtedly affect the accuracy of a “read” since the acids would be crowded together or stretched apart in the blockade time trace affecting the sampling per acid. Second, although the volume differences may not easily be resolved, taken altogether the other components affecting the pore current speculatively make it possible to discriminate between the acids. For example, although the volumes of isoleucine and leucine were practically identical (*V*_I,L_ = 0.166 *nm*^3^, *R_g_*_I_ =0.1833 *nm, R_g_*_L_ =0.1800 *nm*), the blockade current offered the prospect of resolving the two since *I*_L_ = 183.9 ± 42.1 *pA*, *I*_I_ = 137.9 ± 18.4 *pA*, so that Δ*I*_L−I_ = 46 *pA*. On the other hand, despite the volume difference, the largest amino acid residue, tryptophan (*V*_W_ = 0.228 *nm*^3^, *I*_W_ = 131.3 ± 29.3 *pA*, *μ*_W_ = 13.5 ± 0.4 *nm*^2^*/V·μs*) was predicted to be difficult to discriminate from phenylalanine (*V*_F_ = 0.190 *nm*^3^, *I*_F_ = 131.6 ± 22.9 *pA, μ*_F_ = 13.6 ± 0.4 *nm*^2^*/V·μs*) within the error based on the pore current or the acid mobility alone (supplemental Table 1). Likewise, it would be difficult to discriminate cysteine (*I*_C_ = 232.2 ± 3.9 *pA*, *μ*_C_ = 12.9 ± 0.1 *nm*^2^*/V·μs*) from alanine (*I*_A_ = 236.2 ± 11.2 *pA*, *μ*_A_ = 12.5 ± 0.1 *nm*^2^*/V·μs*) based only on the current or mobility. The similarity in the mobility of different acids extracted from these MD simulations may not be so representative, however. There was a threshold velocity below which the peptide fragment did not translocate through the pore at all.

## Calling acids, point mutations and PTMs in proteoforms

In principle, acids with a lower (higher) mobility require a longer (shorter) time to penetrate the waist, stretching (contracting) the position and duration of the *i*^th^ fluctuation, Δ*τ_i_*, in the blockade. To account for dilations or contractions, i.e. Δ*τ*_1_…Δ*τ_i_*#x2026;Δ*τ_n_*, that might result from variations in the acid mobility, the fluctuations in the (normalized) time traces associated with Z-scored empirical consensuses were aligned to the corresponding MD simulations of the pore current using a DTW algorithm (Methods). The DTW algorithm was used to gauge the similarity between unsynchronized features in two time traces of the same length, *n*, and then perform a non-linear warping of the time axis to produce an optimal match between the positions of each feature through a search characterized by a symmetric step pattern through an *n* × *n* matrix containing the Euclidean distances between the points associated with a query and the reference.

As a first test of the algorithm, an empirical query acquired from one *sub*-nanopore (LT111) was aligned to an empirical Aβ_1-42_ reference sequence from another *sub*-nanopore (PM13) to look for differences between the two (Fig. 3). This type of differential analysis was used as a test because it did not appeal to any model, but instead relied only on changes in the blockade current of the query measured relative to the reference to illuminate any differences in the acid sequence. It was compelling evidence of reproducibility that the normalized time sequences of the queries acquired from the same peptide, but with different *sub*-nanopores, were correlated to the reference (*PCC* = 0.32) to begin with (Fig. 3a, red and blue lines, respectively). Strikingly, however, DTW alignment improved the correlation between the two (Fig. 3b) to near perfection (*PCC* = 0.96), although differences in the amplitudes at particular times still remained (Fig. 3f). Other *sub*-nanopores acquiring data from the same peptide performed similarly.

**FIGURE 3.**
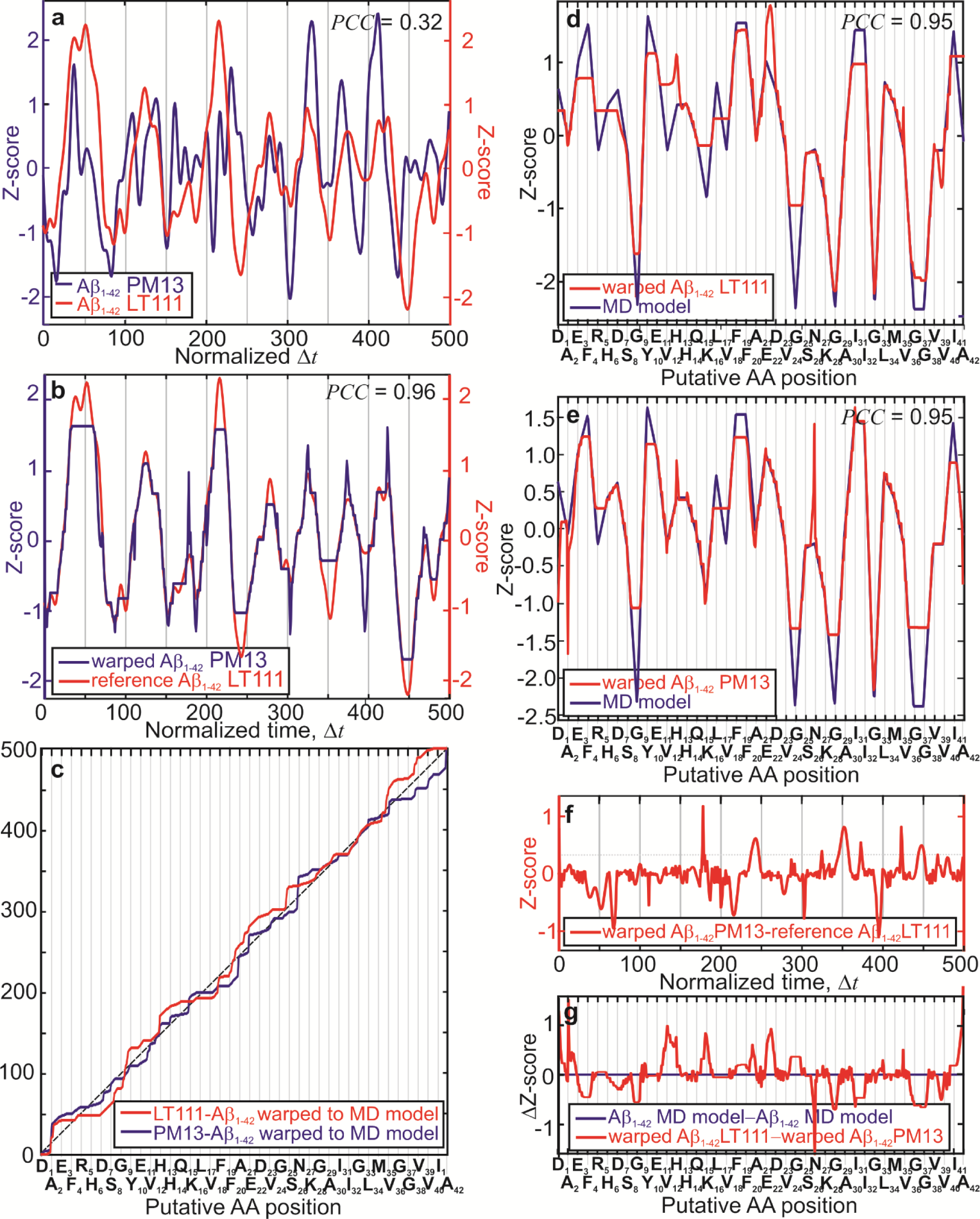
Peptide sequence analysis of Aβ_1-42_ using two different *sub*-nanopores. **(a)** Plots of two Z-scored empirical blockade current consensuses versus normalized duration are juxtaposed. These data were acquired at 0.6 V by forcing denatured Aβ_1-42_ through two different *sub*-nanopores (denoted by PM13 and LT111) with similar topographies at transmembrane bias voltage 0.6 V and measuring the current. Although acquired from different pores, these two traces were correlated (*PCC* = 0.32). **(b)** The Z-scored empirical reference consensus associated with the translocation of Aβ_1-42_ (blue line) through LT111 is shown versus normalized time. To improve the similarity with the reference, juxtaposed on the same plot is shown the <u>d</u>ynamic time-<u>w</u>arped (DTW) version of the Z-scored blockade consensus obtained from PM13 in (a) with a window Δ*τ_i_*, = 2.0 AA wide for DTW. After DTW, the correlation improved to near perfection (*PCC* = 0.96). **(c)** The two empirical consensuses shown in (a) were warped to the reference MD model for Aβ_1-42_ (red and blue lines). The corresponding mappings are shown to illustrate how DTW warps the time-line to improve the similarity. The diagonal (dashed black) line defines a uniform velocity and is used to highlight positional shifts or time delays, Δ*τ_i_*, associated with acid mobility. **(d,e)** Using the mappings in (c), the Z-scored (left vertical axis) empirical consensuses acquired with LT111 and PM13 shown in (a) were aligned to the MD model for Aβ_1-42_. The putative acid positions are shown on the horizontal axis. The aligned empirical data is nearly perfectly correlated to the MD-model (*PCC* = 0.95), but there are “read” discrepancies associated with the smallest and largest acids. **(f)** For reference, the difference between the Z-scores of the two empirical consensus shown in (b), which were aligned to each other, indicating irreproducible noise between different pores. **(g)** The difference between the Z-scores of the empirical consensuses shown in (d,e) each aligned to the Aβ_1-42_ MD model reference is shown versus the putative acid positions (red line). Juxtaposed on the same plot is the difference in Z-scores between the Aβ_1-42_ MD models for each (blue line).

To call the acids from the features in the fluctuation patterns, the empirical consensuses were subsequently aligned to MD simulations of the pore current associated with the translocation of Aβ_1-42_. To align the empirical consensuses to the MD model, the DTW algorithm compensated for local accelerations or decelerations by warping the time axis—shifting an acid-call in normalized time to the left (right) in the map to produce a shorter (longer) Δ*τ_i_* to compensate for a higher (lower) acid velocity in the pore waist. It was evident from the maps linking the queries to the reference model (Fig. 3c, red and blue lines) that both of the Aβ_1-42_ empirical consensuses were indicative of a molecule progressing with a *nearly* uniform velocity through the pore (Fig. 3c, dashed black line), although deviations from a constant velocity were observed frequently. The largest deviations (< 24 samples) from a uniform velocity occurred in both pores near the N-and C-terminus of the peptide, likely due to the frustration associated with threading peptide into the orifice and acceleration when vacating the pore.

Once aligned (Figs. 3d,e), the warped empirical consensuses were observed to be highly correlated with the MD model for Aβ_1-42_ (*PCC* = 0.95 for both). However, regardless of the correlation, the acid glycine with the smallest volume (*V_G_* = 0.0599 *nm*^3^), a practically neutral charge and with the largest number of waters in the waist, (G9, G25, G29, G37 and G38), was routinely overestimated in measurements (< 1.2 σ) relative to the model, but not always (G29 and G33). This deficiency likely represented a lower bound on the dynamic range of the measurement. In support of this assertion, serine (*V_S_* = 0.0917 *nm*^3^) was not usually overestimated. Curiously though, phenylalanine, leucine and isoleucine, with larger volumes (*V_L,I_* = 0.168 *nm*^3^) and a lower number of neighboring waters in the waist compared to glycine, were often (but not always) underestimated in the measurement (F4, F19, F20, L17, I41). The deficiencies near the N- and C-termini might be attributed to irregularities in the molecular velocity and sampling associated with the peptide threading and vacating the pore, but tyrosine (Y10) with the largest volume (*V_Y_* = 0.1912 *nm*^3^) and a low number of waters in the waist was likewise underestimated relative to the prediction of the MD model. So, these deficiencies may instead represent an upper bound on the dynamic range of the measurement.

Three salient aspects were illuminated by this analysis. First, *sub*-nanopores with similar pore topographies produced correlated outcomes. The consensuses representing the same peptide acquired using different *sub*-nanopores (Fig. 3) with similar topographies produced correlated outcomes even without alignment, which indicated a (narrow) fabrication process margin for a *sub*-nanopore that makes large-scale manufacturing possible. Second, the nearly perfect correlation that developed between the MD model and the aligned empirical consensus means that the fluctuations translated to “reads” with single residue resolution,(16) which should facilitate calling the acids as it alleviates the analytical and computational burden associated with the identification of multiple monomers producing a fluctuation in a blockade.(13−15) Finally, despite the nearly perfect correlation, the dynamic range of a “read” measured relative to the MD-model seemed limited at high acid volume (F) when there were fewer waters in the pore, and at the low acid volume (G) when there was more water.

This differential analysis with nearly single acid resolution was used to unambiguously discriminate between Aβ_1-42_ and a scrambled variant, SAβ_1-42_. The empirical consensuses for Aβ_1-42_ and SAβ_1-42_ consisting of 499 and 490 blockades were nearly perfectly aligned (*PCC* = 0.95 for both) to the corresponding MD models with a warping window Δ*τ_i_* = 2.0 AAs (24 samples) wide (Figs. 3b and 4a). However, just as it was in Aβ_1-42,_ glycine (G23, G27, G28 and G37 in SAβ_1-42_), was implicated in the measurement errors (< 1.6σ) relative to the SAβ_1-42_ model (but not always G5 and G13 in SAβ_1-42_). Curiously, while isoleucine (I2) near the N-terminus was underestimated, the acids with larger volumes such as I18, I30, L10, and L36 were accurately appraised in SAβ_1-42_ in contradiction to the results obtained from Aβ_1-42_. Since the same acids were just measured out of sequence, the discrepancies between Aβ_1-42_ and SAβ_1-42_ suggest that neighboring acids may affect a “read”.

**FIGURE 4.**
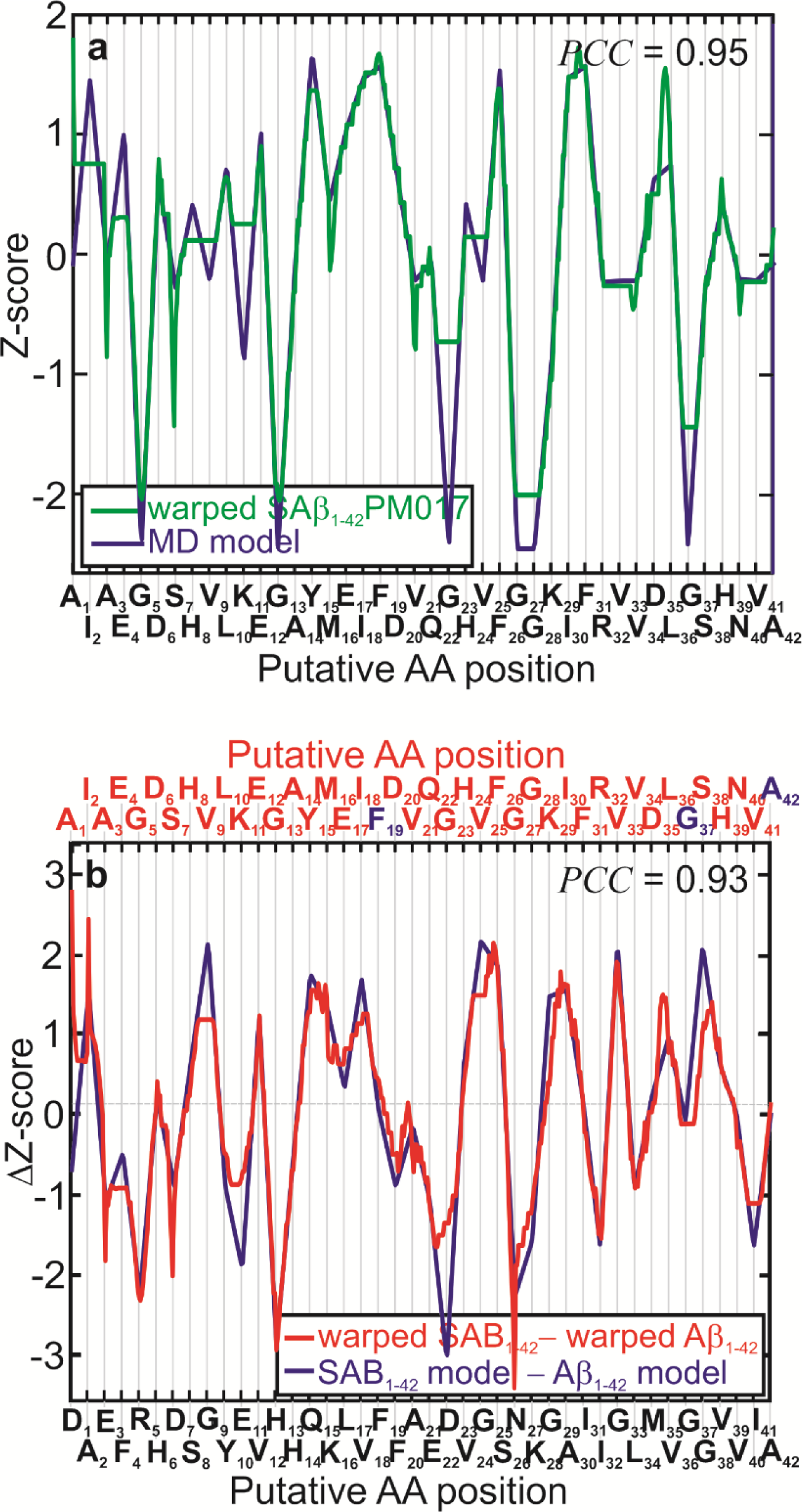
Sequence analysis of beta-amyloid (Aβ_1-42_) and scrambled beta-amyloid (SAβ_1-42_) with single acid resolution using a *sub*-nanopore. **(a)** The aligned empirical consensus for SAβ_1-42_ is shown versus the putative AA positions (red line). The corresponding MD model is juxtaposed on the same plot (blue line). The consensus is well correlated to the model with *PCC* ≥ 0.95. **(b)** The differences in the Z-scores between the aligned consensus for Aβ_1-42_ shown in Fig. 3e and the aligned consensus for the SAβ_1-42_ variant in (a) are shown versus the putative AA positions (red line). Juxtaposed on the same plot is the difference in the Z-scores between the respective MD models for Aβ_1-42_ and the SAβ_1-42_ (blue line). The difference between the empirical consensuses were well correlated to the difference in the corresponding models (*PCC* > 0.93). Based on the close correspondence, the two variants could be discriminated with a *p-value* < 10^-5^.

To highlight the differences and, at the same time, check for consistency, the Z-scores of the aligned empirical data acquired from SAβ_1-42_ and Aβ_1-42_ were subtracted. This type of differential analysis compounds the error, but tests the consistency of the measurement of each acid (relative to the models). Three especially stringent conditions that tested the effect of neighboring acids occurred at sites 19, 37 and 42 where the two peptide sequences shared common acids (F19, G37 and A42). The nearly perfect correlation (*PCC* = 0.93) observed between the aligned empirical difference and the difference in the respective MD−models (Fig. 4b) testifies to the consistency of the measurement, although there were still discrepancies. The largest discrepancies occurred at site 4, implicating phenylalanine (< 0.4 σ), and sites 9, 23 and 38 that implicating glycine (< 1.5σ). On the other hand, the signals attributed to F19, G37 and A42, which were common to both peptides, were accurately read—the same signals were measured with different neighboring acids. This evidence supports the idea that acids in the neighborhood do not always appreciably affect a read. Regardless, signatures were evident in the blockade current fluctuations for both peptides that could be used to identify the sequence of the constituent AA residues. Despite the discrepancies, because the empirical differences so closely tracked the differences in the models, SAβ_1-42_ with the same constituent acids could be discriminated easily from Aβ_1-42_ this way with a *p-value* < 10^-5^.

Predicated on this type of differential analysis, to validate the acid assignments, other proteoforms of Aβ_1-42_ were likewise assayed with a *sub*-nanopore. These proteoforms were selected to test the site specificity, the effect of neighboring acids, the sensitivity to the acid volume and hydropathy. To test site specificity, PTMs of Aβ_1-42_ in which small, hydrophilic S8 and S26 acids were glycosylated with a large, hydrophilic O-linked β-*N*-acetylaglucosamine (denoted as S8OGlcNAc and S26OGlcNAc, respectively), were assayed and compared. Likewise, to test the sensitivity, S26 was phosphorylated (denoted by S26OPO_3_) and then compared with S26OGlcNAc.

Both O-GlcNAcylation and phosphorylation can occur on the same serine residues, which suggests a complex biological relationship between the two PTMs. Like glycosylation (*V*_GlcNAc_ = 0.221 *nm*^3^), phosphorylation also introduces a hydrophilic group in the side chain of serine, but it’s miniscule in size by comparison (*V*_PO4_ = 0.056 *nm*^3^). Correspondingly, MD simulations predicted an increase in the (negative of the) blockade (*I*_S8,26Gly_ = 58.3 ± 28.2 *pA*) relative to serine in the Aβ_1-42_ reference (*I*_S8,26_ = 248.7 ± 45.7 *pA*), but a barely perceptible increase with phosphorylation (*I*_S26PO4_ = 196.7 ± 74.7 *pA*). So, phosphorylation of S26 (i.e. S26OPO_3_) represented an especially stringent assessment of the sensitivity. The Arctic mutation of Aβ_1-42_ in which a very small, hydrophobic G was substituted for a middling volume, hydrophilic E22 (denoted as E22G) was used to gauge the relative importance of volume and hydropathy. Following the same line of argument, it was conjectured that the difference in volumes between E and the G substitution in E22G, (i.e., Δ*V* = *V*_E_ − *V*_G_ = 0.138 − 0.060 nm^3^ = 0.078 nm^3^, supplemental Table 1) would produce a reduction in the blockade current relative to Aβ_1-42_. The MD simulations predicted a correspondingly large difference in the pore current (Δ*I* = *I*_E_ − *I*_G_ = 165.2 ± 19.3 *pA −* 386.9 ± 47.9 *pA = −*221.7 *pA*) and a discrepancy in the number of waters surrounding E (8.2) compared to G (10.2) (Fig. 2d and supplemental Table 1). However, E is charged and hydrophilic, whereas G is neutral and so, it was speculated that the mutation would test the effect of hydration as well as volume since the number of waters displaced from the pore by the acid was also supposed to affect the blockade current.(42) Another measure of the relative importance of volume and hydropathy was the mutant in which the small volume, neutral C was substituted for a very large volume, hydrophobic F at site 20 (denoted by F20C). MD predicted (Δ*I* = *I*_F_ − *I*_C_ = 131.5 ± 22.9 *pA −* 232.2 ± 3.9 *pA = −*100.7 *pA*) for this mutation. Finally, the Flemish mutation, F4W, provided the ultimate test of the MD-model in which a neutral acid with the largest volume, W, was substituted for another large volume acid that was hydrophobic, F. MD simulations predicted practically no difference in the pore current (Δ*I = I*_W_ − *I*_F_ = 131.3 ± 29.3 *pA* − 131.6 ± 22.9 *pA =* 0.3 *pA*).

The empirical consensuses for these variants were aligned to the corresponding MD-models using DTW with a warping window Δ*τ_i_* ≤ 2.0 AA (24 samples) wide (Figs. 5a,c,e,g and supplemental Figs. 4a,c). Generally, the aligned empirical data was tightly correlated to the respective MD-models (*PCC* ≥ 0.90) with the usual exceptions. The pore current assigned to glycine (G9, G25, G29, G33, G37 and G38) was routinely, but not always, overestimated relative to the model, whereas the current attributed to isoleucine (I31, I32, I41), leucine (L17) and phenylalanine (F19, F20) were (sometimes) underestimated. To highlight the differences, the Z-scores of the aligned empirical data, acquired from each of the variants and Aβ_1-42_, were subtracted. Commensurate with the predictions, the difference in the Z-scores for S26OGlcNAc and S8OGlcNAc revealed prominent (> 2.1σ) peaks at S26 and S8 (Figs. 5b,d) that were attributed to glycosylation there. Although a prominent peak (1.7σ) was observed in the difference between the Z-scores associated with the aligned S26OPO3 and the Aβ_1-42_ consensuses at S26 as well (Fig. 5h, red line), that identification was equivocal. Consistent with the simulations, a nadir in the Z-score (3.5σ) of the empirical consensus for the E22G mutation relative to the Aβ_1-42_ reference was evident at site 22 (Fig. 5h, red line). Curiously, the measurement of the fluctuation amplitude attributed to G22 in the mutant was assessed perfectly relative to the MD-model, as was G25. This was curious because measurements of all the other variants routinely overestimated G25 relative to the MD-model, but not in E22G. Based on this evidence, it was conjectured that acids three sites removed might influence a “read” and that even the smallest acid could provisionally be “read” accurately, depending on the sequence.

**FIGURE 5.**
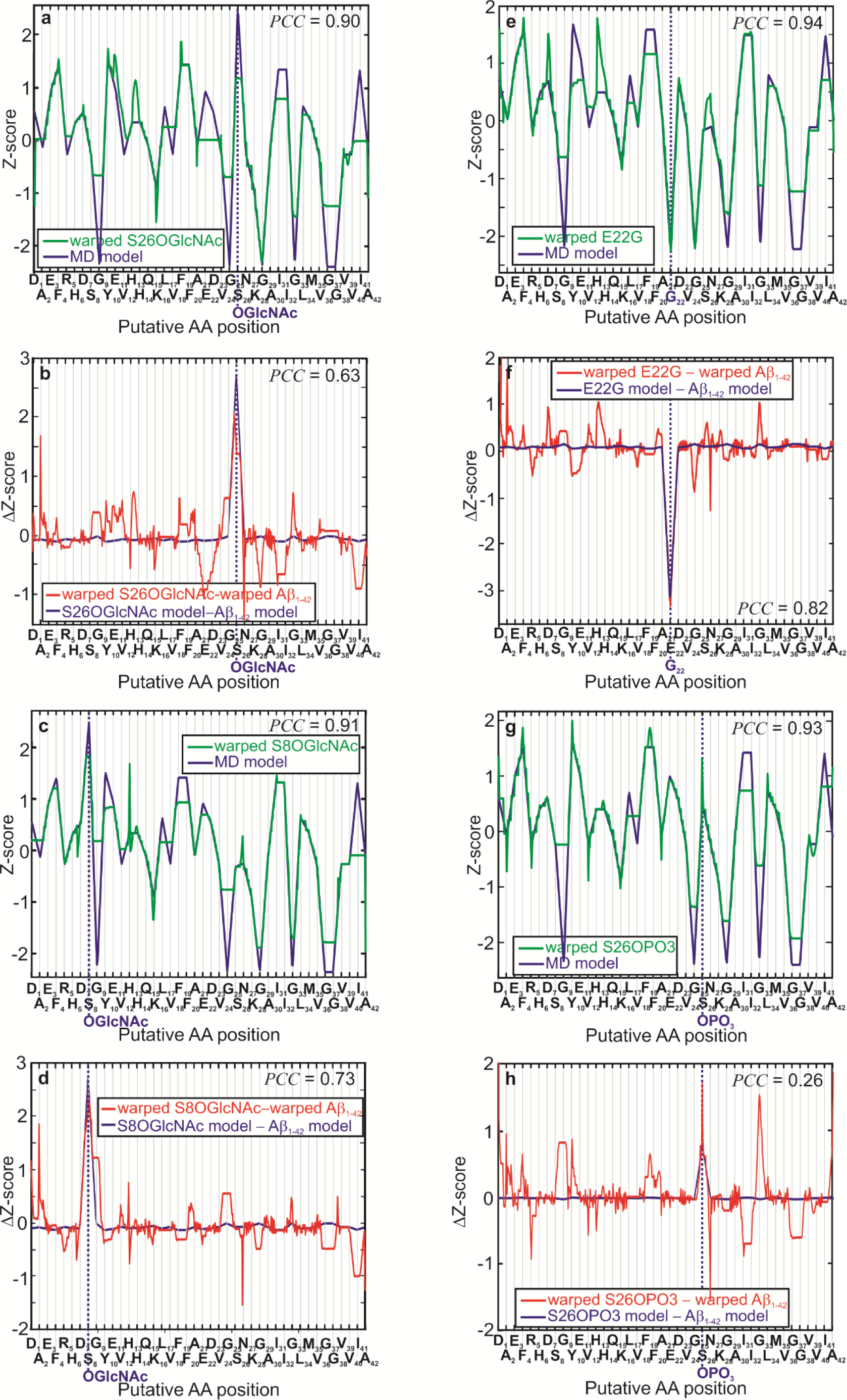
Proteoform sequence analysis with near single acid resolution using a *sub*-nanopore. **(a,c,e,g)** Empirical consensuses associated with blockades due to S26GlyNAc, S8GlyNAc, E22G and S26OPO3 are shown after alignment to the corresponding MD models versus the putative AA positions (red lines). Juxtaposed with the empirical consensuses are the corresponding MD models for each variant (blue lines). The dotted blue line indicates the site of the mutation or PTM. The aligned consensuses were all well correlated to the respective MD models with *PCC* ≥ 0.90. **(b,d,f,h)**The differences in the Z-scores between the aligned consensus for Aβ_1-42_ and the respective aligned consensuses for the variants in (a,c,e,g) (red lines) are shown versus the putative AA positions. (The Aβ_1-42_ reference data used for comparison—see Fig. 3e—which was acquired with the same *sub*-nanopore, was nearly perfectly aligned with the respective MD model *PCC* = 0.95 as well.) Juxtaposed on the same plots are the differences in the Z-scores between the respective MD models for Aβ_1-42_ and the corresponding variants (blue lines). The difference between the empirical consensuses were correlated to the difference in the corresponding models (0.26 < *PCC* < 0.82). The peaks prominent in (b,d) were attributed to the glycosylation S26 and S8. The nadir evident in (f) was attributed to the E22G substitution.

The accurate “reading” of the miniscule G prompted a chase to the bottom of the *sub*-nanopore sensitivity using the two mutants F20C and F4W (supplemental Fig. 4). A nadir was barely perceptible at site 20 in the difference in Z-scores for the aligned consensus acquired from the F20C mutant relative to the Aβ_1-42_ reference (supplemental Fig. 5b), but the identification with the mutation was equivocal. Likewise, the difference between the F4W mutant and Aβ_1-42_ consensuses at site 4 was not perceptible (supplemental Fig. 4d), but not expected according to MD, which predicted that Δ*I = I*_W_ − *I*_F_ = 0.3 *pA*.

Taken altogether including the controls, these data aligned to MD simulations proved that measurements of fluctuations in the blockade current through a *sub*-nanopore when a proteoform is impelled through it were sensitive enough to “read” the acids in the primary structure and identify point mutations and PTMs in them precisely with single site specificity. The alignment exposed irregularities in the timing of the fluctuations, however, which were attributed to variations in the acid mobility in the pore in line with MD simulations. Finally, the acid sequences were inferred from clusters formed from a few hundred blockades, which indicated that only a minimal amount of material was required to identify the peptide.

## Methods/Experimental

### Peptide Synthesis

The eleven carrier-free peptides used in these experiments were either purchased directly or custom-synthesized (Anaspec, Fremont, CA; Biosynthesis, Lewisville, TX) and then reconstituted according to the protocols offered by the manufacturers. (The constituent acid residues comprising the beta-amyloid variants included nineteen of the twenty proteinogenic AAs; only proline was not included, although it was included in the controls.) To avoid aggregation and false readings, the peptides were reconstituted in a 5 nM solution with PBS without adding BSA. Typically, 20 µL protein solution from a 5 nM aliquot was mixed with 50 µL of 0.1% sodium dodecyl sulfate (SDS) stock; 50 µL out of 10 mM β-mercaptoethanol (β-ME) stock; 120 µL of 500 mM *NaCl* electrolyte and 760 µL of 250 mM *NaCl* electrolyte to produce a 100 pM protein solution containing 0.005% (v/v) SDS and 500 μM β-ME in 1 mL of 250 mM *NaCl* buffer in a 1.5 mL centrifuge tube. The temperature of the tube was raised to 85°C for 2 hours to denature the protein and then the solution was cooled (to 23°C) and injected into the *cis*-reservoir of the <u>p</u>oly<u>d</u>i<u>m</u>ethyl<u>s</u>iloxane (PDMS) microfluidic device bound to the silicon chip supporting the membrane with a pore through it. The *trans*-reservoir usually contained about 1 mL of a solution consisting of 900 μL of 250 mM *NaCl* electrolyte with 50 μL of 550 mM *NaCl* electrolyte along with 50 μL of 0.1 % (v/v) SDS.

### *Sub*-nanopore fabrication and visualization

A *sub*-nanopore sputtered through a thin <u>a</u>morphous <u>si</u>licon (*a*-*Si*) membrane was used to analyze the peptides. The native oxide that forms on amorphous silicon(31–34) motivated the choice of this membrane because: 1. it was apparently self-limiting and likely stabilized the pore topography; 2. it supposedly compensated for silicon surface states and thus, minimized the charge in the pore; 3. it was supposed to be negatively charged in water, which precluded bio-fouling; and 4. due to the silanol (*Si-OH*) on the surface, it was hydrophilic, which should ostensibly facilitate frictionless sliding translocation kinetics through the pore.(14)

Custom-made *a*-*Si* membranes nominally ranging from 5-8 nm thick were manufactured (SiMPore, Inc. West Henrietta, NY) by sputter-depositing silicon on a 60 nm thick G-FLAT® *SiO*_2_ layer on a float-zone silicon handle 100 μm thick and subsequently, sandwiched with another *SiO*_2_ layer with the same 60 nm thickness followed by the deposition of 150 nm of tetra<u>e</u>thyl <u>o</u>rtho<u>s</u>ilicate (*TEOS*). The *TEOS* was densified to decrease porosity and improve its performance as an etch mask. The *a*-*Si* film was sputter-deposited on a rotating substrate 25° off normal in 5-30 mTorr *Ar* gas with a large substrate-target spacing (10 cm) and low (150°C) substrate temperature. The deposition rate was about ∼0.04 nm/s. This was supposed to produce an *a*-*Si* film with minimal dangling-bond density and incorporated gas, but in one lot (#2331), to increase the density and reduce leakage current due to asperities the substrate temperature was increased from 150°C to 200°C during the deposition of the *a-Si*. From this sandwich, membranes < 4-25 μm on-edge were revealed using an aqueous solution of <u>e</u>thylene <u>d</u>iamine and <u>p</u>yrocatechol (*EDP*) chemical etch of the silicon through a silicon nitride window defined by photolithography on the polished backside of the handle wafer and then a 10:1 <u>b</u>uffered <u>o</u>xide <u>e</u>tch (*BOE*) was used to remove the oxide. Although large area (25 μm square) thin *a*-*Si* membranes were fabricated reproducibly with acceptable yield, they were fragile so the area of the membrane had to be minuscule to provide a margin for manufacturing. So, membranes only a few micrometers on-edge were manufactured by controlling the size of the nitride window on the backside of the handle wafer (Fig. 1a).

Finally, the membrane thickness was important because it supposedly affected the field distribution in the pore and therefore the resolution of a “read” (supplemental note #1 and Fig. 1). Thin membranes that can support a large electric field were required to lessen the prospect for backward steps in the motion of the acid through the pore to facilitate the interpretation of the blockade current. Generally, the manufacturer’s (SiMPore’s) specifications were relied upon for an estimate of the membrane thickness, but as a check the thicknesses of select *a*-*Si* membranes were also measured directly by transmission <u>e</u>lectron <u>m</u>icroscopy (TEM) (Fig. 1b) or <u>s</u>canning <u>e</u>lectron <u>m</u>icroscopy (SEM) by sacrificing a sister chip from the same lot fabricated similarly or by using <u>E</u>lectron <u>E</u>nergy <u>L</u>oss <u>S</u>pectroscopy (EELS). Although EELS indicated that the membranes were thin, in our hands it proved to be an inconsistent measure of the thickness and so, electron microscopy was used to accurately gauge the thickness. The thickness was measured by puncturing the membrane or cleaving through the *Si* chip and inspecting an up-turned edge. Generally, the thicknesses were observed in the range from *t* = 3.5 to 6.0 nm in accord with the SiMPore specifications. Finally, the roughness of the top surface of the membrane, measured with custom-built silicon cantilevers (Bruker, Fremont, CA) with 2 nm radius tips, was estimated to be < 0.5 nm-rms.

Just prior to loading it into the STEM column, the membranes were plasma cleaned using Tergeo-EM (PIE Scientific, Union City, CA USA). The Tergeo-EM was operated at 10 W using an 80% Ar+20% O_2_ gas feed in a down-stream, pulse mode (1/16 duty-cycle, which was cycled twice for a total exposure of 2 min) such that the samples were actually outside the plasma (to eliminate sputtering) and subjected to only extremely short plasma pulses (to reduce the intensity). Subsequently, pores were sputtered through the thin *a*-*Si* membrane using a tightly focused, high-energy (300 keV) electron beam carrying a current ranging from 60−460 pA (post-alignment) in a (<u>S</u>canning) <u>T</u>ransmission <u>E</u>lectron <u>M</u>icroscope (STEM, FEI Titan 80-300 or aberration-(probe) corrected FEI Themis Z, or Spectra 300, ThermoFisher, Hillsboro, OR) with a <u>F</u>ield <u>E</u>mission <u>G</u>un (FEG). The sputtering time varied depending on the microscope conditions and the membrane. The sputtering was either timed or directly tracked by monitoring the Ronchigram until a pore was observed—typically, a pore opened in < 90 sec (supplemental Fig. 6), but sometimes exceeded 3 min depending on the microscope focus conditions and the lot.

A Ronchigram is just another name for a convergent beam diffraction pattern focused on an amorphous thin film. The center of it has high local magnification and is supposed to be a practically aberration-free portion of the beam. To sputter a *sub*-nanopore using a Ronchigram as a guide, the convergence point of the probe is exactly focused onto the specimen, at which point the magnification of the Ronchigram becomes infinite and the intensity becomes uniform. Thus, by adjusting the Ronchigram to have a uniform intensity, the exact focus of the incident beam on the specimen can be obtained. A measurement of the angular extent of the uniform intensity then determines the angular range of the incident probe with no aberration, which is characterized by the semi-angle, *α*. In an (aberration) probe-corrected STEM, the <u>d</u>epth <u>o</u>f field (DOF) or focal depth, Δ, scales like: Δ ≍ *λ* /*α* ^2^ (= 6.1 *nm*@300 *kV* ). (43) So, the vertical extension of the probe decreases with increasing *α* and decreasing *λ*. A shallow probe like this can be used for sputtering, but only on precisely focus.

After sputtering, the pore was sometimes re-acquired with either <u>H</u>igh <u>R</u>esolution <u>T</u>ransmission <u>E</u>lectron <u>M</u>icroscopy (HRTEM) or <u>H</u>igh-<u>A</u>ngle <u>A</u>nnular <u>D</u>ark <u>F</u>ield (HAADF−)STEM. To minimize beam damage, the pores were examined using low beam current (< 16−30 pA) or low voltage (80 kV) or both. The illumination convergence angle in the Titan was typically *α* = 10 mrad at 300 kV, whereas in the Themis Z and Spectra 300, *α* > 18−32 mrad at 300 kV or *α* = 27.1 mrad at 80 kV with a monochromator limiting the energy dispersion in the range 200−220 mV at 80 kV according to EELS. Whereas the Themis Z and Spectra STEM resolution at 300 kV were determined to be < 60 pm on a *GaN* lattice, the resolution at 80 kV was < 120 pm, according to a dumbbell lattice image acquired from (110) crystalline silicon. Regardless, the high-resolution, aberration-corrected probe used in HAADF-STEM facilitated the direct interpretation of the images in terms of the mass density under the beam without resorting to multiple views or simulations.

The pores sputtered through the *a*-*Si* membrane were likely oxidized in the ambient, which might account for the change in the topography observed in Figs. 1c,d. Surprisingly, oxidation did not completely obstruct a pore this size, possibly due to stress near the waist. It has been shown that the stress from deformation during silicon oxidation retards the reaction rate and (oxygen) diffusivity, and that the oxidation rate is generally retarded for sharp curvatures, especially at low temperature.(31−34)

### Microfluidics

The silicon chip supporting the membrane with a single pore through it was bonded to a PDMS (Sylgard 184, Dow Corning) microfluidic device formed using a mold-casting technique; all of which has been described previously.(16,44) The PDMS microfluidic was formed from a thoroughly stirred a (15:1) mixture of elastomer (siloxane) with a curing agent (cross-linker) cast in a mold formed from DSM Somos ProtoTherm 12120 plastic (Fineline Prototyping, Raleigh, NC) and then degassed and cross-linked at 75°C for 2 hrs. The microfluidic device consisted of a microchannel (each 250 × 75 μm^2^ in cross-section) on the *trans*-side connected by a *via* that could be as small as 75 μm in diameter to a well on the *cis*-side (supplemental Fig. 5a).

To reduce the parasitic capacitance, either a thick layer of photo-sensitive polyimide (κ = 2.8-3.5) was spin-deposited onto the *Si*-chip and then patterned with UV-lithography using the membrane as a mask to produce a *via* with a > 25 μm diameter prior to sputtering a pore or a thick layer of <u>p</u>oly<u>d</u>i<u>m</u>ethyl<u>s</u>iloxane (PDMS, κ = 2.3−2.8) with a >75 μm diameter *via* that was bound to the silicon chip after sputtering a pore. To apply the polyimide, first, an adhesion promotor (Pyralin, VM-652) was spun on top of the chip for 30 sec., and then on top of that polyimide photoresist (HD8820, HD MicroSystems) was deposited at 10,000 rpm to produce a film thickness of 5.2 ± 0.6 μm. After a soft-bake at 120°C for 3 min, a window the size of the membrane was opened in the resist with a 3 min UV flood exposure, followed by development in AZ 917 MIF developer and rinsed in 18 MΩ-cm <u>d</u>e-ionized (DI) water. Subsequently, the polyimide layer was hard-baked at 250°C/300°C for 3-6 h to remove all solvents. Although the noise reduction was appreciable (supplemental Figs. 5b,c), the yield from chips with a polyimide coating was low. So, most of the work was accomplished using a PDMS laminate to mitigate noise. The PDMS laminate with a *via* over the membrane mitigated current noise also, but the lower κ required a thicker layer. While a noise reduction with *vias* as small as 75 μm in diameter was observed (supplemental Figs. 5d,e), the main effect on the noise was captured without an aperture by the overlap of the narrow *trans*-channel on the *Si*-chip.(18,44)

Generally, a tight seal was formed between the silicon chip containing the *a*-*Si* membrane with the pore in it and the PDMS microfluidic channel with a plasma-bonding process. First, the *trans*-side was plasma-bonded to a clean 75 × 25 mm^2^ glass slide, 1 mm-thick (VWR, Radnor, PA) using a (blue-white) 25 W oxygen (1200 sccm *O*_2_) plasma (PDS-001, Harrick Plasma, Ithaca, NY) for 30 sec. Then, to promote adhesion and produce a hydrophilic surface while eliminating sputtering, the *Si* chip was held in vacuum, but outside the plasma, and subjected to short plasma pulses. This was accomplished using a down-stream remote plasma at 1/16 duty cycle produced in a Tergeo-EM plasma cleaner (PIE Scientific, Union City, CA USA). The Tergeo-EM was operated at 10 W with a gas feed of 10 sccm *O*_2_ and 3 sccm *Ar*. At the same time, the PDMS microfluidic device was exposed to an oxygen (800 sccm) plasma in Harrick for 30 sec and subsequently, within 1 min the two were brought into contact under an inverted optical microscope (Axiovert-40, Zeiss) using a micromanipulator with a vacuum probe to precisely position the *Si* chip in the PDMS microfluidic. To ensure a >100 GΩ seal to the PDMS, the back (*cis*) side of the silicon chip was painted with (15:1) PDMS, and then the ensemble was heat-treated at a temperature of 85°C for 24 hr in vacuum. Subsequently, *Ag/AgCl* electrodes (Warner Instruments, Hamden, CT) were embedded in each channel to independently, electrically address the *cis-* and *trans*-sides of the membrane. Supposedly, the impedance at high frequency (1 MHz) diminishes, which motivated the choice of *Ag/AgCl* electrodes.(45) The two microfluidic channels were connected to external pressure and fluid reservoirs through polyethylene tubing at the input and output ports.

Finally, to minimize electrical noise, the microfluidic chip was encased in a die-cast aluminum NEMA enclosure with coaxial electrical feedthroughs that contacted the *Ag/AgCl* electrodes immersed in the *cis*- and *trans*-reservoirs. The assays of denatured peptides were performed at 4−7°C to minimize aggregation and slow the translocation velocity.(46) This was accomplished by first refrigerating the whole assembly, i.e., the chip, microfluidic device and metallic case, to 4°C (for about 20 min) and then sandwiching it between two ice blocks for the duration of the electrical measurements.

### Low-noise electrical measurements

Prior to the actual measurement, the *sub*-nanopore was wetted by first exposing the device to a remote oxygen 20 sccm, 20 W, 1/16 duty-cycle for 30 sec in the Tergeo-EM plasma cleaner (PIE Scientific, Union City, CA USA), and then immediately after exposure to plasma (within 1 min), the *cis*- and *trans*-channels in the microfluidic device were immersed in 250 mM *NaCl* electrolyte. Instead of a plasma, methyl alcohol was sometimes used to wet a *sub*-nanopore. With this option, the microfluidic device with a *sub*-nanopore embedded in it was placed in a vacuum desiccator alongside a Petri dish filled with methyl alcohol. Subsequently, the desiccator was held in partial vacuum for about 5 hrs after which time the *cis*-reservoir and *trans*-channel in the microfluidic device were immersed in 250 mM *NaCl* electrolyte. To remove air bubbles, the microfluidic device was also subjected to a vacuum for < 10 min and then “electro-wetted” by sweeping the transmembrane voltage over the range extending from ±0.6 V in 60 sec intervals (using the Axon “pCLAMP 10.7.0 software, Molecular Devices, San Jose, CA) while monitoring the pore current. The *sub*-nanopore was successfully “wet” when a featureless current trace (without blockades) was observed, which amounts to performing effectively a negative control for each *sub*-nanopore. Generally, absent air bubbles, no blockades were observed beyond the noise in controls that comprised just the electrolyte after electro-wetting.

Subsequently, to measure the blockade current with protein in the *cis*-reservoir, a transmembrane voltage bias (< 0.70 V) was applied to the reservoir relative to ground in the *trans*-channel using *Ag/AgCl* electrodes and the corresponding pore current was measured at 4−7°C. To minimize drift and reduce acoustic noise, all the measurements were performed on an optical air table with active piezoelectric vibration control (STACIS, TMC, Peabody, MA), housed in an acoustically isolated, NC−25 (Noise criterion) rated room in which the temperature was stabilized to less than ± 0.1°C over 24 hr through radiative cooling.

The low bandwidth electrical measurements followed a procedure similar to that described elsewhere.(16) Tersely, a transmembrane voltage (< 0.70 V) was applied to the reservoir (containing 75 µL of electrolytic solution and 75 μL of 2× concentrated solution of protein and denaturant) relative to the ground in the channel using *Ag/AgCl* electrodes and the corresponding pore current was measured at 5 ± 0.1°C using either an Axopatch 700B or an Axopatch 200B amplifier (Molecular Devices, San Jose, CA) with an open bandwidth. The actual bandwidth was inferred from the rise-time to a sharp (10 ps rise-time) input pulse to be about 50 kHz to 100 kHz, depending on the amplifier and the feedback. The analog data were digitized by a 16-bit DigiData 1440A or 1550B data acquisition system (DAQ, Molecular Devices, San Jose, CA) at a sampling rate of 250-500 kS/sec and recorded in 3-minute-long acquisition windows and stored in <u>a</u>xon <u>b</u>inary files (ABFs).

To improve the “read” accuracy, the *sub*-nanopore current was amplified with a wide-band (1.8 MHz) trans-impedance (current) amplifier (e.g., DHPCA-100, FEMTO) sampled with a DAQ at 3.6 MS/sec (PCIe-6374, NI, Austin, TX). Typically, the data were recorded continuously and filed every 15 sec (55 MS per file). If a pore fouled, to rehabilitate it, the microfluidic device was flushed with 18 MΩ−cm DI water for 1 da. To establish that the pore was cleared, the open pore current and current noise were measured again prior to an experiment. The pore was reused only if the noise and the open pore current returned to that observed in the pristine pore. If this procedure failed to clear it, 0.1% SDS solution was flushed through the channel to disperse latent aggregated protein in an attempt to recover the pore.

### Signal recovery and reconstruction

The blockade current distribution was important for accurate signal reconstruction. The fractional change in current has been linked to the ratio of the molecular volume to the pore volume according to: Δ*I* / *I*_0_ = *f* · Δ*V_mol_* / *V_pore_* · *S*, where *f* measures the molecular shape and orientation, and *S* is a size factor that accounts for distortions in the electric field that occur when the molecule is comparable in size to the pore.(13, 38) For the average acid volume constituting Aβ_1-42_ (Δ*V_AA_* = 0.13 nm^3^) occluding the high electrical field full <u>w</u>idth at <u>h</u>alf-<u>m</u>aximum (FWHM) stretch (0.7 nm long) in a bi-conical (10°) *sub*-nanopore with a mean diameter of 0.40 nm at the waist, it was estimated that Δ*V*_1*AA*_ / *V_pore_* = 0.13 / 0.118 ≍ 1 ∝ Δ*I* / *I*_0_, which overestimates the observed median fractional blockade. (Coincidently, on the other hand, if current crowding was ignored, and the molecular volume contained within the entire bi-conical pore (30°) spanning a 5 nm membrane is taken into account, then thirteen acids span the membrane so that: Δ*V*_13 *AA*_ / *V_pore_* = 1.7 /16.1 = 0.11 ∝ Δ*I* / *I*_0_ .) Thus, based on these estimates, it was concluded that the empirical mean fractional blockade over this range was consistent with a single molecule in the pore.

To inform on the sequence, ideally each acid should be sampled at least twice, at the Nyquist rate, but if the acid velocity through the pore was variable then the sampling would not be uniform in time. The width of the blockade duration distribution relative to the median provides a hint about the molecular velocity variation. Based on the median duration of the translocation, the average velocity through the pore was about *v̄* = 2.3×10^−4^ *nm* / *ns*(= 0.23 *mm* / *sec*). The force on an Aβ_1-42_ molecule for a bias voltage of 0.70 V was measured directly to be (on average) about *F̄* = 25 *pN* as it slid relatively frictionless through a *sub*-nanopore (supplemental Fig. 2). (14) The Einstein relation between the ionic mobility (*μ*) and the diffusion constant (*D*) predicts that: *D* = *k_B_Tμ* / *q*. = *k_B_Tv̄*^2^ / *F* = 5.7 ×10^−14^ *m*^2^ / *sec* and so, assuming each acid moving with a mean velocity of a Gaussian profile, the ratio between the duration of a translocation and the FWHM of the distribution is given by: 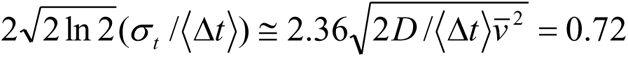. Thus, the width of the blockade duration distribution should be about 72% of the median, which corresponds closely to the width of the red contour shown (Fig. 1g) that represents 50% of the total population. In other words, Brownian diffusion accounts for the width of the distribution of the blockade duration.

The velocity fluctuations inherent in Brownian diffusion could confound a “read” if backward fluctuations in the acid motion through the pore exceed the distance between acids (i.e., if the molecule back-steps). According to the Nyquist criterion, the minimum bandwidth required to “read” each acid moving with a mean velocity of *v̄* = 2.3×10^−4^ *nm* / *ns* is about 1.2 MS/sec. To avoid a back-step and a mistaken “read” then, the contributions to the root-mean-square thermal velocity fluctuations (up to 1.2 MHz), 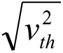, must be smaller than the average acid velocity or, in other words, 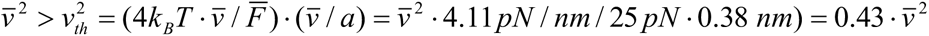. Thus, a “read” error was tied to the ratio of the thermal energy to the work accomplished impelling an acid through the pore,(39) which is about 4*k_B_T* · *v̄*/(*F̄a*) < 0.4 in these measurements. So, to avoid “read” errors due to back-stepping, a large electric force, *F*, was indicated, which translated to a short median duration Δ*t* and a wide bandwidth amplifier for detection.

Signal recovery and reconstruction were accomplished using customized code written in MATLAB and/or Python. There were five steps in the process: **1.** identification of blockades from a digitally filtered version of the empirical current traces; **2.** selection of a workable subset from the unfiltered empirical data and then uniformly sampling and re-scaling each event; **3.** digitally filtering and identifying those blockades with similar fluctuation patterns to produce tight clusters; **4.** identifying reverse-time clusters using the <u>d</u>ynamic time <u>w</u>arping algorithm (DTW) distance between clusters; and finally, **5.** aligning the empirical consensus with a MD model using a DTW algorithm.

**1.** First, events were detected from empirical current traces subjected to a low-pass, 8-pole Bessel filter with a 20 kHz cutoff frequency using *OpenNanopore* (47) with an *S* = 3.8 deviation from the baseline and end at *E* = 0 from the baseline with a filter coefficient *F* = 0.999 and 10 baseline points included for the CUSUM−fit. Events that qualified as blockades satisfied the criteria that the open pore current *I*_0_ ≥ 150 pA with *I*_0_ after the event was within 10 pA of the open pore current prior to the event. The events that qualified as blockades were subsequently extracted from the unfiltered empirical data, along with a header and footer 200 samples long, which extended beyond the duration of the blockade so that any artifacts associated with filtering would be divorced from the signal. The blockade, Δ*I*, the (local) open pore current, *I*_0_, and blockade duration, Δ*t*, were calculated for each event using data filtered through an 8-pole low pass Bessel filter. Typically, the number of blockades detected in a single experiment ranged between 2000 and 5000.
**2.** A subset of the empirical data was selected by culling blockades with a duration too short to satisfy the Nyquist criterion or too long to preclude back-stepping or slip-and-stick motion.(14) Likewise, blockades with a fractional blockade current, Δ*I/I*_0_, beyond the 50% contour (Fig. 1g, red highlight) were also culled. Three subsets were compiled by zero-phase-shift filtering each blockade with a 5^th^ order low-pass Butterworth IIR filter with a pass-band defined by the 3dB point at (the low frequency) 17-cycles/(½-blockade duration) and (the high frequency) 42-cycles/(½-blockade duration), and with a 5^th^ order pass band Butterworth IIR filter defined by the 3dB points at 1.5-cycles/(½-blockade duration) and 42-cycles/(½blockade duration). The 200-point headers and footers were removed after filtering. Since the durations were variable, the number of samples in each blockade also varied.(48−50) So, each of the selected blockades were re-sampled to a normalized timescale with 500 samples. Finally, since the blockade current varied (within 10% of *I*_0_), the mean current was zeroed, and the result normalized by the standard deviation (*σ* ) to form the Z-score or the standard score for each blockade.
**3.** The selected, renormalized blockades were subsequently clustered using an <u>a</u>ffinity <u>p</u>ropagation (AP) algorithm implemented in MATLAB (apcluster.m) recursively. First, a subset filtered at 17 cycles was used to compute the blockade cross-correlation distance matrix. Based on a distance matrix, AP grouped blockades into clusters. Indices of clustered blockades were then used to assemble consensuses from the subset filtered at 42 cycles. The cross-correlation distance matrix was computed for clustered consensuses and passed to the AP algorithm. AP grouped similar clusters into a super-cluster.
**4.** To form the final consensus with AP, super-clusters with the shortest mean Euclidean distance between the filtered elements or the shortest median duration were selected. A second super-cluster was then identified by the shortest DTW distance to the time-reversed version of the first. The elements of the latter super-cluster were time-reversed and combined into a final consensus. The resulting consensus terminals were trimmed (typically by 5 points) and then resampled to 500 points.
**5.** Finally, DTW was used to align the time-axis of the empirical consensus to a model of the peptide in which each acid in the acid sequence was represented by the current derived from MD simulations. The DTW algorithm implemented in Python can be found on-line at: https://github.com/statefb/dtwalign.(20) DTW computes the local stretch or compression to apply to the time axes of two time-series in order to optimally map one (query) onto the other (reference). DTW efficiently and accurately accomplished the alignment along the time axis, but it suffered limitations. For example, depending on the number of samples and the width of the warping window, DTW created so-called “singularities” when the amplitudes of the features in different time sequences varied.(16,22) These “singularities” developed when the algorithm tried to offset variability in the amplitude by warping the time-axis, which sometimes produced one-to-many mappings in which a single point in one time series was mapped to an entire sub-section of another. The endpoints were also important and their definition was uncertain likely because of irregular sampling as the peptide threaded or vacated the pore. Due to the boundary conditions that required the warping path to start and finish in diagonally opposite corners, DTW could be invariant to all warping, but still could not handle the difference at the start (or end) of the two time-series. This constraint meant that the algorithm was forced to match the pairs of beginning and end points, which adversely affected the statistical distance measure. So, to circumvent large variability in the endpoints, the algorithm used to identify a blockade was implemented with an extraction criterion. The endpoints were chosen to maximize the *PCC* MD model; the uncertainty in the definition of the endpoints was always less than 24 samples and typically < 5 samples. Finally, to avoid overfitting, each consensus was constrained to 500 samples and the warping window was restricted to a range of Δ*τ_i_* ≤ 2.0 AA positions or 24 samples out of 500.

### Finite Element Simulations (FES)

The FESs were performed by using COMSOL (v5.6, COMSOL Inc., Palo Alto, CA, USA) as described elsewhere.(17) Tersely, the FESs were based on continuum modeling, which accounted for a bi-conical shape of the particular pore, the reduced electrophoretic mobility and the steric effect of ions explicitly. The electrohydrodynamics were governed by coupled Poisson and Stokes equations. The applied potential φ and the potential *ψ* due to charges in the pore were de-coupled from one another and solved independently. The relationship between *ϕ* and the charge carriers, e.g., *Na^+^* and *Cl^−^*, is given by the Poisson equation, ∇^2^*ψ* = −*ρ* / *εε*_0_, where *ρ*, *ε* and *ε*_0_ were the volume charge density and the relative and vacuum permittivities, respectively. The charge density is given by *ρ* = *F* ∑_*i*_*z_i_c_i_* where *F* = 96,485 C mol^−1^ is the Faraday constant, *z_i_* is the valence and *c_i_* is the molar concentrations of *i*^th^ ionic species in the bulk. Electro-osmotic flow was captured by the Navier–Stokes equation: i.e., *η*∇^2^*u* − ∇*p* − *F* ∑_*i*_*z_i_c_i_* ∇*V* = 0, where the total potential, *V* = φ + *ψ*, *η* is the viscosity, *p* is the pressure and *u* is the velocity vector. The transport of ionic species is described by the Nernst–Planck equation given by: *D*_*i*_∇^2^*c_i_* + *z_i_μ_i_c_i_*∇^2^*V* = *u* ·∇*c_i_*, where *D_i_* is the diffusion coefficient and *μ*_i_ is the ionic mobility of the *i*^th^ species. Thus, *u, V* and *c_i_* are coupled between equations. The boundary conditions are specified in the Supplemental Table 3 and material properties such as the diffusivity in the *sub*-nanopore were restricted by the literature as elaborated elsewhere.(17)

Following earlier work,(17) to estimate the pore conductance the radial distributions of the electric potentials and ion concentrations were calculated, assuming that the radius-dependent concentration followed a Boltzmann distribution according to: ∇^2^Ψ = sinh(Ψ) / *λ*^2^ (1+*α*(cosh(Ψ) −1)), Ψ = *eψ* / *k*_B_*T* where *ψ* is the (radius-dependent electric potential), *k*_B_ is the Boltzmann constant; *T* is the absolute temperature; and *l* is the Debye screening length and *α* = 2*a*^3^*n*, where *a* is the radius of the solvated ion. The topography of each pore was taken into account to match the data. With the assumption of a pore topography, the effective thickness of the membrane was estimated from the electric field distribution using FESs. With this estimate, the surface charge density was then inferred from measurements of the pore conductance at dilute concentration, assuming initially that the cations carried the current prodominately and negligible electro-osmotic flow.

Parenthetically, to account for the current crowding near the waist of the pore (supplemental Fig. 1c), a convolution was performed over the AA volumes using a window *k*−acids wide that usually spanned between 1 ≤ *k* ≤ 2 AAs, in correspondence with FESs of the electric field distribution. Then the width of the window was optimized to maximize the *PCC* between the model and the empirical data. However, convolutions with 1 ≤ *k* ≤ 2 offered only a modest improvement in the *PCC* so, for clarity, the results presented here were compared to a *k* = 1 model exclusively, although it does not necessarily represent the optimum.

### <u>M</u>olecular <u>D</u>ynamics (MD) Simulations

For an accurate assessment with atomic detail, following earlier work,(51−53) the ion and peptide transport was simulated in a *sub*-nanopore through a silica membrane by MD. All simulations were performed using NAMD version 2.14 in NVT ensemble.(52) To construct the *sub*-nanopore, initially, a crystalline silica slab was generated with a number of unit cells required to fill the pore-excluded volume of the amorphous silica (*SiO*_2_) membrane with a density of 3.0 g/cm^3^. The surface geometry of a double conical *sub*-nanopore having an elliptical cross-section was modeled using the relation:

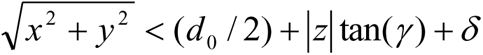

where *γ* = 10° is the cone angle at the minimum cross-section of the *sub*-nanopore near the waist, *d*_0_ = 0.4 *nm* is the diameter of the pore at the waist and *δ* is the van der Waals radii of the pore surface atoms (taken as 0.2 nm). The *Si* and *O* atoms were randomly placed inside the membrane, excluding the volume of the *sub*-nanopore, and then the entire membrane was subjected to simulated annealing at a high temperature 7000 K. A repulsive grid potential was applied to avoid the entry of *SiO*_2_ atoms inside the *sub*-nanopore and CHARMM force field parameters were used for *Si* and *O* atoms in the membrane.(54) Later, the bonds between the *Si* and *O* atoms of silica were defined using a 0.22 nm distance cut-off. Using this protocol, a silica *sub*-nanopore was created like the actual topography (Fig. 1d). A similar protocol was successively used to obtain the stable silica pores in previous studies.(53−56) In the process of creating the pore, the resulting total charge of the membrane was typically non-zero so, to maintain the electro-neutrality, the partial charges on the *Si* atoms were adjusted so that the total charge of the membrane was zero.

Although the net charge on the membrane is zero, due to the amorphous structure of the annealed *SiO*_2_ used, some small charges at the surface of the pore wall due to partial charges on *Si* and *O* were expected. We define the atoms on the surface of the pore wall as those that lie within 0.8 nm of the surface of the pore wall, and it is only the charges on these atoms that influence water, ions, and peptide. The number of atoms which satisfy this criterion is around 2400 and when the net surface charge on the pore due to these atoms is evaluated, a small, net negative charge (−6*e*) on the surface is observed over the 2400 atoms. The net effect of this small negative charge does not cause any counter-ions to accumulate on the surface as these are distributed across the surface.

The MD simulations were performed for the β-amyloid peptide and fragments of it translocating through the pore spanning the membrane. The coordinates for *β*-amyloid (pdbcode:1IYT) peptide were obtained from the <u>p</u>rotein <u>d</u>ata <u>b</u>ank (PDB);(57) peptide fragments of it were constructed from VMD using CHARMM36(59) topologies and force-fields. The glycosylated serine variant with O-linked *β*-N-acetylaglucosamine (O-GlcNAc) was constructed using CHARMM-GUI.(58) The constructed peptide structures were placed at the orifice of the pore to study their translocations through the entire silica *sub*-nanopore and CHARMM36 force field parameters were employed for the peptide fragments.(59) The constructed silica *sub*-nanopore-protein complex was solvated with a 5 nm thick TIP3P water film on either side of the pore to screen the electrostatic interactions of the pore with its periodic images.(60) The *Na*^+^ and *Cl*^−^ counter ions were added to the solvated pore to set 200 mM *NaCl* concentration. Initial systems were subjected to 2000 steps of conjugate-gradient minimization to remove the bad contacts and followed by 1 ns NPT equilibration at temperature 300 K with a 1 fs time step for integration to achieve the bulk water density of 1.0 g/cm^3^. Periodic boundary conditions were used in all three directions.

A Nosé–Hoover thermostat with a damping coefficient of 1 ps was connected to silica atoms of the membrane to maintain the temperature.(61) Subsequently, the equilibrated systems were subjected to a 20 ns NVT MD simulation with a 0.12 nm cutoff for calculating the short-range part of the non-bonding interactions and the long-range electrostatic interactions were calculated using <u>P</u>article-<u>M</u>esh-<u>E</u>wald (PME) method.(62)

A 5 V potential drop was applied across the membrane spanned by the *sub*-nanopore system for each of the peptide fragments to induce the ionic flux and subsequently, the fragments were impelled through the pore by pulling with a zero-acceleration velocity of 5×10^-6^ *nm/ps*. The ionic current inside the *sub*-nanopore was computed using:

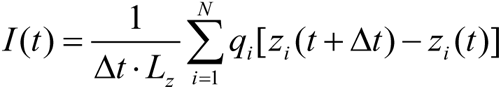

where *z_i_* and *q_i_* were the *z*-coordinate and atomic charge of the *i^th^* ion, respectively. The parameter *N* represents the total number of ions present in the pore and *L*_z_ represents the *z*-dimension of the system (14 nm). The pore current, AA mobility, hydration, charge and gyration radii, *R_g_*, at the minimum cross-section of the pore were computed using the peptide translocation trajectories and then the data for the entire *β*-amyloid peptide and its variants, E22G and S26GlcNAc, were constructed and arranged according to the pre-determined sequence of β-amyloid from PDB. An acid was taken to be within the pore waist when −0.3 nm < *z* < 0.3 nm, with the origin at 0, which demarcates the region of interest. All the data, including the current, were collected when the fragments were in this *z*-range.

As mentioned above, single-acids with a small zero-acceleration velocity of 5×10^-6^ nm/ps were used to estimate the currents associated with the translocation of the amino acids, after which the current profile for the whole sequence was constructed. The velocity used to pull the acids was adjusted so that it takes about 3-4 ns to translocate through the pore waist. In addition, averaged values of current are estimated from 5 simulations for each amino acid and the standard errors associated with them were calculated. This ensured that the currents obtained for each acid converged and the errors associated with the MD simulations were minimized, which facilitated a comparison to the empirical results since enough statistics were acquired to capture the physics effectively.

### Force and Pore Current Molecular Spectroscopy with an <u>A</u>tomic <u>F</u>orce <u>M</u>icroscope (AFM)

The force and current data were acquired from single protein molecules using a customized AFM (MFP-3D-BIO, Asylum Research, Santa Barbara, CA) interfaced to an inverted optical microscope (Axio-Observer Z1, Zeiss), all enclosed within a Faraday cage, as described elsewhere.(14) Tersely, a low noise Z-sensor in the AFM was coupled with an ultra-quiet Z-drive to produce noise in the tip-sample distance < 30 pm at 1 kHz bandwidth. To minimize drift and reduce acoustic noise, an inverted optical microscope was mounted on an optical air table with active piezoelectric vibration control (STACIS, TMC, Peabody, MA), housed in an acoustically isolated, NC-25 (Noise criterion) rated room in which the temperature was stabilized to less than ±0.1 °C over 24 h through radiative cooling. Temperature fluctuations appeared to be a source of long-term drift but with temperature regulation, the drift of the system was reduced to 0.6 nm/min. Sound couples strongly into the microscope and was another potential source of instrument noise. Therefore, acoustically loud devices, especially those with cooling fans such as power supplies, amplifiers, and computers, were placed outside of the room. With these precautions, force detector noise is < 10 pm/√Hz for frequencies above 1 Hz; the on-surface positional noise measured < 45 pm A-dev.

To acquire the data, first, the topography of the membrane and the location of the pore relative to the edges of the membrane were determined in air in non-contact (tapping) mode using a silicon cantilever (SSS-FM, Nanosensors, Neuchatel, Switzerland) with a 2 nm nominal radius, and a spring constant ranging from *K_spr_* = 0.5-9.5 nN/nm and a 45-115 kHz resonant frequency (in air). Force spectroscopy was performed in a 250 mM *NaCl* electrolytic solution, using a custom MSNL silicon cantilever (Bruker, Camarillo, CA) without metal reflex with a 2 nm tip radius. Each MSNL probe includes six cantilevers with a range of force constants (10 < *K_spr_* < 100 pN/nm) and resonant frequencies (7< ω_0_ <125 kHz). Considering only the off-resonance thermal noise of the cantilever 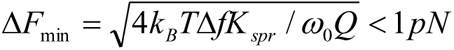 where typically *K_spr_* ∼ 10-30 pN/nm, *Δf* = 100-3300 Hz was the measurement bandwidth, *ω*_0_ = 2π×5.3 -18 kHz was the angular resonance frequency of the cantilever, and *Q* ∼15−17(0.8−1.3) was the quality factor in air (liquid).

To locate the pore relative to fiducial marks, i.e., the edges of the (4 × 6 μm^2^) membrane, an AFM topographical scan was performed with a sharp tip in liquid in constant force (contact) mode. After that, the pore location was re-acquired in liquid with a second cantilever on the same probe through triangulation from the fiducial marks and a small area scan. Then a constant +0.70 V was applied to the anode on the *trans*-side of the membrane with the cathode on the *cis*-side grounded, and the pore current was measured continuously with an external amplifier, while the force on the cantilever was determined from the deflection. Starting from a position about 100-120 nm above the membrane, the tethered protein, immersed in a solution of 250 mM *NaCl* electrolyte and SDS, was repeatedly advanced towards the sub-nanopore at 20 nm/s, captured and threaded through it by the electric field, and then retracted from it at a constant 4 nm/s velocity by the AFM while the current, tip deflection and *z*-position were recorded. The tip position above the membrane was determined from the sum of the tip deflection and Z-sensor position.

The ionic current through a nanopore was measured in a Faraday cage with a patch-clamp amplifier (Axopatch 200B, Molecular Devices, San Jose, CA) in whole-cell mode. *Ag/AgCl* electrodes embedded in the microfluidic device were used to establish a transmembrane potential. An electrical bias of +0.70 V was applied between *Ag/AgCl* electrodes embedded in the *trans*- and *cis-*channels, respectively, and the current between them was measured using an Axopatch 200B amplifier at 17.6 ± 0.1 °C. Each data channel was subsequently digitally filtered at 5 kHz and sampled at 10 kHz and then digitally filtered again using a 100 Hz eight-pole Bessel filter (MATLAB, 2016a).

## ACKNOWLEDGEMENTS

This work was mainly supported by a grant from the Open Philanthropy Project, and partially supported by the Keough-Hesburgh Professorship. Some of the aberration-corrected STEM was accomplished at the Materials Research Laboratory Central Research Facilities at the University of Illinois in Urbana, IL with the assistance of J. Mabon and C.Q. Chen. Some of the data was acquired and preliminary analysis accomplished by Tautvydas Lisauskas (Max Planck Institute) and Juan Oviedo Robles (Intel). We gratefully acknowledge discussions with Xiaowen Liu (Tulane University), Jun Li (Notre Dame) and Winston Timp (Johns Hopkins University).

## DATA AVAILABILITY

Summaries of the data generated and/or analyzed during the current study are included in the published article and the corresponding supplemental information file. Requests for these materials should be addressed to GT.

## Supporting Information

The supporting information includes:

1. Supplemental Table 1 showing <u>a</u>mino <u>a</u>cid (AA) residue volumes used in the peptide analysis.
2. Supplemental Table 2 showing the sequences of amyloid-beta and variants of it.
3. Supplemental Note #1 describing the calculation of the electric field in a *sub*-nanopore.
4. Supplemental Table 3 showing the COMSOL simulation parameters.
5. Supplemental Figure 1 showing <u>F</u>inite <u>E</u>lement <u>S</u>imulation (FES) of a *sub*-nanopore.
6. Supplemental Figure 2 showing the forces and blockade current measured as a single Aβ_1-42_ peptide was impelled relatively frictionless through a *sub*-nanopore.
7. Supplemental Note #2 explaining random electrical current noise in a *sub*-nanopore.
8. Supplemental Figure 3 illustrating a low-bandwidth test of the sensitivity of a *sub*-nanopore to molecular volume using single block co-polymers and a consensus of Aβ_1-42_ blockades that is used to suppress random noise.
9. Supplemental Figure 4 showing proteoform sequence analysis with near single acid resolution using a *sub*-nanopore.
10. Supplemental Figure 5 illustrating noise mitigation with a *sub*-nanopore through an *a-Si* membrane laminated with thick polyimide and PDMS.
11. Supplemental Figure 6 illustrating how a Ronchigram is used to monitor the sputtering of a pore with a tightly focused, high energy electron beam in a <u>S</u>canning <u>T</u>ransmission <u>E</u>lectron <u>M</u>icroscope (STEM).
12. Supplemental VideoAB1-42e shows a MD simulation visualizing the translocation of beta-amyloid through a *sub*-nanopore.

