## Supplementary Materials for "Decoding Proteoforms with Single Acid Resolution Using a *Sub*-nanometer Diameter Pore"

### SUPPLEMENTAL MATERIALS for
### De-coding Proteoforms with Single Acid Resolution Using a *Sub*-nanometer Diameter Pore

*Apurba Paul,[1] Archith Rayabharam,[2] Punam Murkute,[1] Lisa Almonte,[1] Eveline Rigo,[1] Zhuxin Dong,[1] Ashutosh Kumar,[1] Joshy Joseph,[1] Narayana Aluru,[2] and Gregory Timp[1]**

[1]Electrical Engineering and Biological Science, University of Notre Dame, Notre Dame, IN 46556 USA

[2]Department of Mechanical Engineering, University of Texas, Austin, TX 78712-1229 USA

[#]contributed equally

**SUPPLEMENTAL TABLE 1.** Amino acid (AA) residue volumes used in the peptide analysis. The AA and glucosamine volumes were taken from the averages reported in: [P] Perkins, S.J. Protein volumes and hydration effects. *Eur. J. Biochem.* **157,** 169–180 (1986). These averages were generally consistent with those in: [Z] Zamyatnin, A.A. Protein volume in solution. *Prog. Biophys. Mol. Biol.*, **24**,107-123 (1972); and [T] Tsia, J., Taylor, R., Chothia, C. & Gerstein, M. The Packing Density in Proteins: Standard Radii and Volumes, *J. Mol. Biol.*, **290,** 253-266 (1999). The hydropathy ranking 1-20 was taken from [B] Bonella, S., Raimondo, D., Milanetti, E., Tramontano, A. & Ciccotti, G. Mapping the Hydropathy of Amino Acids Based on Their Local Solvation Structure. *J. Phys. Chem. B.* **118,** 6604-6613 (2014). For comparison, the hydropathy indices is also shown from [K] Kyte, J. & Doolittle, R.F. A simple method for displaying the hydropathic character of a protein. *J. Mol. Biol.* **157,** 105-132 (1982).

| **Amino acid** | **Abbreviation** | **Molecular mass (Da)** | **# of atoms** | **MD pore current (pA)** | **Volume (nm$^3$)** [P]/[Z]/[T] | **Hydropathy** [B]/[K] |
|---|---|---|---|---|---|---|
| Alanine | A | 89 | 13 | 236.2 ± 11.2 | 0.0878/0.0886/0.0901 | 13/1.6 |
| Arginine | R | 174 | 26 | 245.8 ± 30.5 | 0.1882/0.1734/0.1928 | 3/-12.3 |
| Asparagine | N | 132 | 17 | 244.4 ± 6.8 | 0.1201/0.1141/0.1275 | 1/-4.8 |
| Aspartic acid | D | 133 | 16 | 191.5 ± 24.1 | 0.1154/0.1111/0.1171 | 8/-9.2 |
| Cysteine | C | 121 | 14 | 232.2 ± 3.9 | 0.1054/0.1085/0.1035 | 12/2 |
| Glutamine | Q | 146 | 20 | 234.9 ± 47.0 | 0.1451/0.1438/0.1494 | 7/-4.1 |
| Glutamic Acid | E | 147 | 19 | 165.2 ± 19.3 | 0.1409/0.1384/0.1408 | 6/-8.2 |
| Glycine | G | 75 | 10 | 386.9 ± 47.9 | 0.0599/0.0601/0.0638 | 14/1 |
| Histidine | H | 155 | 20 | 204.5 ± 35.2 | 0.1563/0.1532/0.1593 | 2/-3 |
| Isoleucine | I | 131 | 22 | 137.9 ± 18.5 | 0.1661/0.1667/0.1649 | 19/3.1 |
| Leucine | L | 131 | 22 | 183.9 ± 42.1 | 0.168/0.1667/0.1646 | 20/2.8 |
| Lysine | K | 146 | 24 | 289.0 ± 4.5 | 0.1727/0.1686/0.1700 | 4/-8.8 |
| Methionine | M | 149 | 20 | 203.6 ± 42.6 | 0.1652/0.1629/0.1677 | 16/3.4 |
| Phenylalanine | F | 165 | 23 | 131.6 ± 22.9 | 0.1897/0.1899/0.1935 | 15/3.7 |
| Proline | P | 115 | 17 | 178.7 ± 23.2 | 0.1233/0.1127/0.1231 | 17/-0.2 |
| Serine | S | 105 | 14 | 248.7 ± 45.7 | 0.0917/0.089/0.0942 | 5/0.6 |
| Threonine | T | 119 | 17 | 251.8 ± 53.8 | 0.1183/0.1161/0.1200 | 9/1.2 |
| Tryptophan | W | 204 | 27 | 131.3 ± 29.3 | 0.2279/0.2278/0.2317 | 11/1.2 |
| Tyrosine | Y | 181 | 24 | 124.2 ± 9.9 | 0.1912/0.1936/0.1971 | 10/-0.7 |
| Valine | V | 117 | 19 | 245.2 ± 12.0 | 0.1388/0.140/0.1391 | 18/2.6 |
| water | $H_2O$ | 18 | 3 | | 0.0245/0.0297 | |
| $PO_4^{3-}$ | $PO_4^{3-}$ | 95/79.97 | 5 | | 0.0565 | |
| glucosamine | GlcNAc | 221.21 | 30 | | 0.222.0±0.002 | |

**SUPPLEMENTAL TABLE 2.** Sequences of amyloid-beta ($A\beta_{1-42}$) and variants of it. The blue symbols indicate the substitutions/mutations sites in $A\beta_{1-42}$ whereas the red letters indicate the acids sites in common with $A\beta_{1-42}$.

| | |
|---|---|
| **$A\beta_{1-42}$** | *DAEFRHDSGYEVHHQKLVFFAEDVGSNKGAIIGLMVGGVVIA* |
| **$SA\beta_{1-42}$** | *AIAEGDSHVLKEGAYMEI**F**DVQGHVF**G**GKIFRVVDLGSHNV**A*** |
| **S26OGlcNAc** | *DAEFRHDSGYEVHHQKLVFFAEDVG-**S(OGlcNAc)**-NKGAIIGLMVGGVVIA* |
| **S8OGlcNAc** | *DAEFRHD-**S(OGlcNAc)**-GYEVHHQKLVFFAEDVGSNKGAIIGLMVGGVVIA* |
| **E22G** | *DAEFRHDSGYEVHHQKLVFFA**G**DVGSNKGAIIGLMVGGVVIA* |
| **S26OPO3** | *DAEFRHDSGYEVHHQKLVFFAEDVG-**S(O-PO$_3$)-**NKGAIIGLMVGGVVIA* |
| **F4W** | *DAE**W**RHDSGYEVHHQKLVFFAEDVGSNKGAIIGLMVGGVVIA* |
| **F20C** | *DAEFRHDSGYEVHHQKLVF**C**AEDVGSNKGAIIGLMVGGVVIA* |
| ***block-p*(R)$_{10}$-*co*-(G$_3$S)$_3$-G** | *RRRRRRRRRRGGGSGGGSGGGSG* |
| ***block-p*(K)$_{10}$-*co*-(G$_3$S)$_3$-G** | *KKKKKKKKKKGGGSGGGSGGGSG* |
| ***block-p*(K)$_{24}$-*co*-(A$_8$P$_8$)3-A$_8$** | *KKKKKKKKKKKKKKKKKKKKKKKKAAAAAAAAPPPPPPPP AAAAAAAAPPPPPPPPAAAAAAAA* |

**SUPPLEMENTAL NOTE #1: The electric field and current in a *sub*-nanopore.**

The precision with which *sub*-nanometer-diameter pores were fabricated translated directly to very stringent control of the electric field and current distribution. Finite element simulations (FESs) of the electric field and the electro-osmotic flow were performed using COMSOL (v5.6, COMSOL Inc., Palo Alto, CA), following a modified Poisson-Boltzmann formalism described in: Rigo, E., et al. Measurements of the Size and Correlations between Ions using an Electrolytic Point Contact. *Nat. Commun.,* **10,** 2382 (2019).

For the simulations, a conformal layer of amorphous silicon dioxide ($SiO_2$) membrane covering an amorphous silicon (*a-Si*) membrane was constructed with a bi-conical pore through it with a *sub*-nanometer-diameter waist. The cone angle ranged from 5–15° near the waist, but expanded to 25-40° near the orifice of the pore. It was assumed that the pore was immersed in 250 mM *NaCl* and a transmembrane bias voltage was applied across the membrane. The boundary conditions for the system are given in supplemental Table 3. The thin (3.5–6 nm) membranes, in conjunction with the bi-conical topography, tightly focused the electric field along the pore axis as evident in supplemental Fig. 1, for $L_m$ = 5 nm with a bias voltage of 0.6 V. Thus, the potential (and the electric force) in a pore was confined to a 0.70 nm region near the constriction (full width-half maximum), which corresponds to about 1–2 AAs in a linear peptide chain assuming that the spacing between $\alpha$-carbons is about 0.38 nm (supplemental Fig. 1b). Support for these assessments of the potential was obtained from measurements of the open pore current-voltage characteristics of *sub*-nanopores. Typical current-voltage characteristics are shown in supplemental Fig. 1. They were approximately linear over a range of ± 1V. A line-fit to this data yielded a conductance of 330 pS for a 0.4 nm-diameter in 250 M *NaCl*. The concentration dependence of the conductivity suggests that there is a fixed charge in the pore (in this case σ = –0.05 $e/nm^2$). Accounting for the surface charge, finite element simulations (FES) can reproduce quantitatively the measured conductance (red dashed line, supplemental Fig. 1c).

**SUPPLEMENTAL TABLE 3:** Finite Element Simulation Parameters.

| Feature | Boundary Conditions |
|---|---|
| 1. *sub*-nanopore | $d$ = 0.4 *nm* diameter, $L_m$ = 5 *nm thick* 10° bi-conical structure |
| 2. *a-Si* membrane | $t_{Si}$ = 5 *nm*; $\varepsilon_r$ = 11.7 |
| 3. $SiO_2$ membrane | $t_{ox}$ = 1 *nm* thick, σ = –0.05 $e/nm^2$; $\varepsilon_r$ = 3.9 |
| 4. Electrolyte | 250 mM *NaCl*; $\varepsilon_r$=78.5; $\eta$ = 1.002 mPa·s; $D_{Na}$ =0.07×1.33×$10^{-9}$ $m^2/s$; $D_{Cl}$ = 0.007×2.03×$10^{-9}$ $m^2/s$. |
| 5. zeta-potential, $\zeta$; ionic radius, $a$, and $\alpha$ | $\zeta$ = –0.007 $V$; $a$ = 0.15 nm ; $\alpha$ = 0.0010162 |

**SUPPLEMENTAL FIGURE 1.**

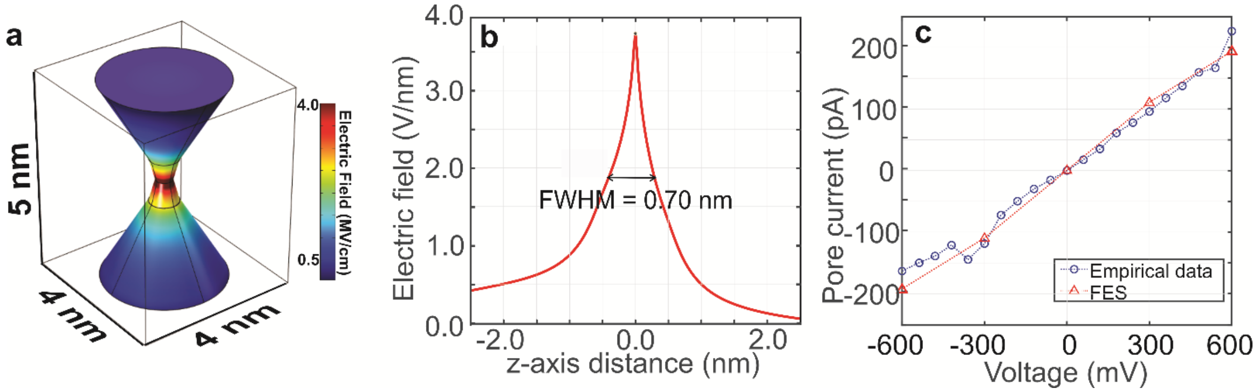


**SUPPLEMENTAL FIGURE 1. Finite Element Simulation (FES) of a *sub*-nanopore. (a)** Based on the *sub*-nanopore topography inferred from HAADF-STEM after oxidation, a three-dimensional (3D) model was used to estimate the electric field distribution along the pore (*z*-) axis. What's shown is a schematic representation of a *sub*-nanopore through a 5 nm thick membrane with a bi-conical topography that is 0.4 nm diameter at the waist with a 10° cone-angle that opens to 30°. **(b)** An FES of the electric field distribution in the pore is shown. The full-width at half-maximum (FWHM) of the electric field distribution is 0.70 nm. **(c)** A measurement of the current-voltage characteristics of an actual *sub*-nanopore immersed in 250 mM *NaCl* electrolyte (blue circles) is shown along with FES simulations of it.

**SUPPLEMENTAL FIGURE 2.**

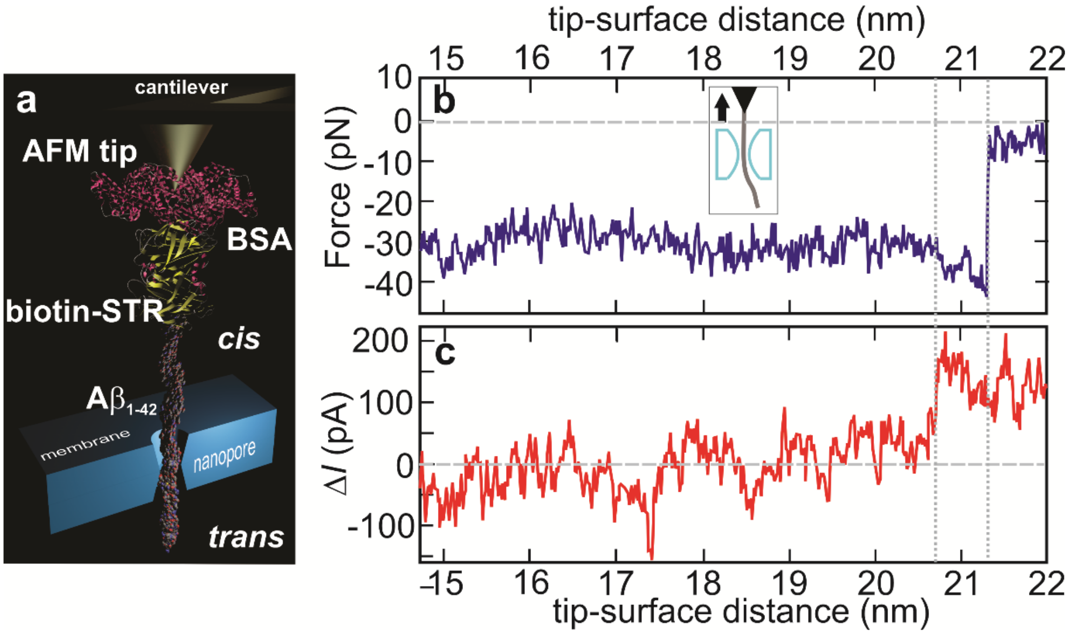


**SUPPLEMENTAL FIGURE 2. The forces and blockade current measured as a single A$\beta_{1\text{-}42}$ peptide was impelled relatively frictionless through a *sub*-nanopore.** **(a)** A schematic cutaway of the experiment showing a biotinylated A$\beta_{1\text{-}42}$ peptide tethered to the tip of an AFM cantilever through a bond to streptavidin (STR), translocating through the *sub*-nanopore. A denatured, tethered peptide-SDS aggregate was advanced by the AFM toward the *sub*-nanopore, captured and threaded through it by the electric field, and then retracted from it at a constant velocity, while the tip deflection and current were recorded to measure the force and blockade current. **(b,c)** Concomitant measurements of the force and current are shown as the A$\beta_{1\text{-}42}$ peptide was impelled through a *sub*-nanopore with a 0.8 × 0.9 nm$^2$ cross-section at the waist spanning an *a-Si* membrane nominally 5 nm thick. Both the *cis*- and *trans*-sides of the membrane were immersed in 0.01% (w/v) SDS in a 250 mM *NaCl* solution. (The *sub*-nanopore cross-section was measured *in situ* immediately after sputtering, but the actual cross-section was likely reduced after exposure to the ambient.) Changes in both the force (b) and the pore current (c) were observed while the AFM cantilever was retracted from the pore at velocity of 2.0 nm/s. The change in the force (b) on the protein that is observed near 21.3 nm is an indication that the electric force on the molecule associated with an applied potential of +0.70 V was terminated. The change in the pore current (c) observed near 20.7 nm indicates the position where the blockade in the pore current was relieved. Only a small force (24 ± 5 pN) was required to extract the molecule from the *sub*-nanopore and relieve the $\Delta I$ = 130 ±40 pA current blockade. The force hardly varied over the distance of the extraction and so it is referred to as "frictionless". **(b, inset)** The cartoon shows the assumed molecular configuration with the arrow indicating the direction of the cantilever motion.

**SUPPLEMENTAL NOTE #2: Random electrical current noise in a *sub*-nanopore.**

The random current noise associated with a *sub*-nanopore assay can be analyzed into four components: thermal, 1/*f*, dielectric and amplifier noise (supplemental Figs. 3d,e).(18) The thermal noise power spectral density (*PSD*) that is associated with the pore resistance was negligible over the frequency band < 2 MHz. Likewise, 1/*f* –noise was negligible above 100 kHz. On the other hand, above 100 kHz the dielectric noise associated with parasitic membrane capacitance and amplifier noise predominated.(17,18) It was observed that the rms-current noise in an open pore, which is equivalent to $\sigma$ when the mean current is suppressed, averaged over durations comparable to a blockade, diminished as $1/\sqrt{N}$ where *N* denotes the number of events with a duration indicative of random noise (supplemental Fig. 3e). Unlike the random noise, however, the fluctuations in the blockade current persisted even after signal averaging with an amplitude swinging between 5–6$\sigma$ (Figs. 1h,i and supplemental Fig. 3f). Thus, the combination of filtering and clustering blockades improved the signal-to-noise ratio.

**SUPPLEMENTAL FIGURE 3.**

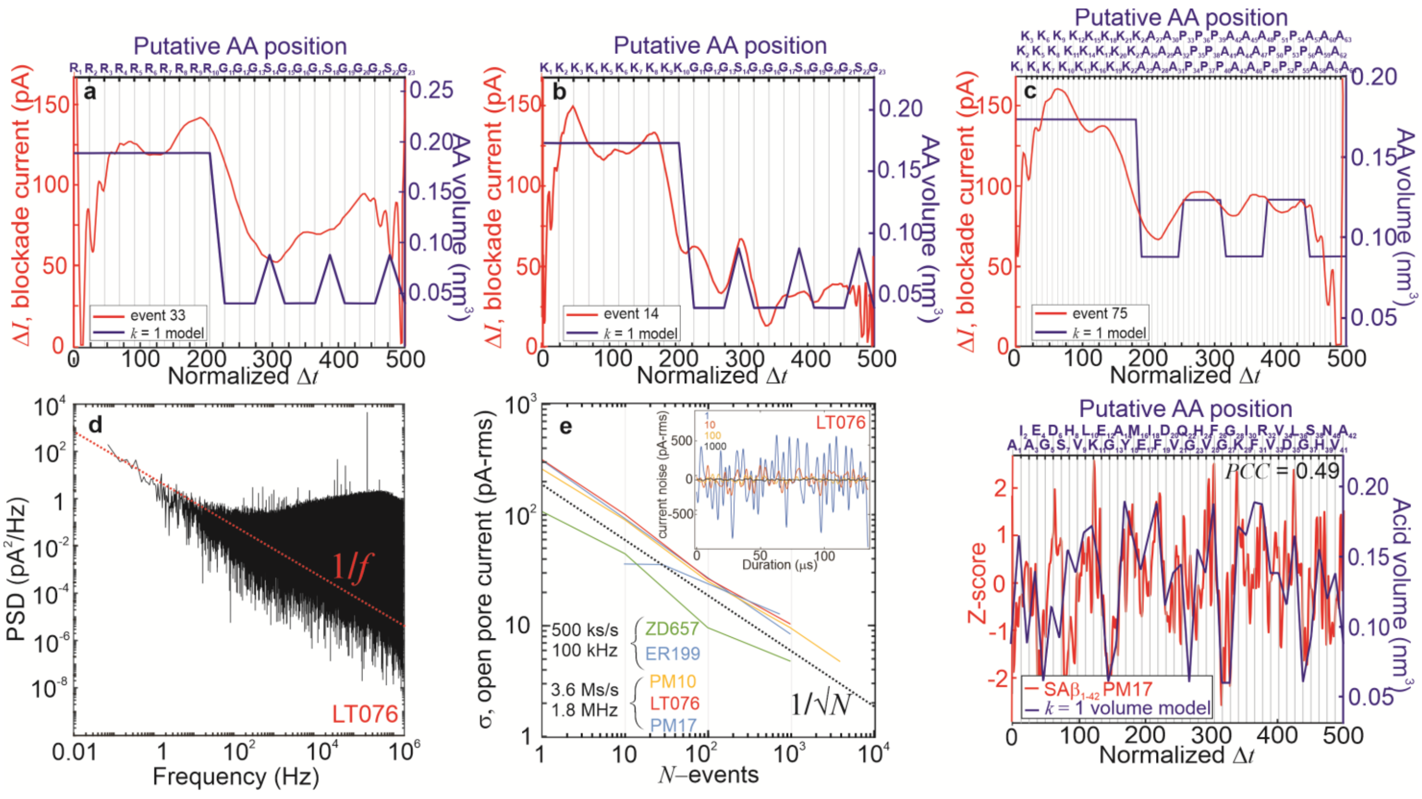

**SUPPLEMENTAL FIGURE 3. Testing the sensitivity of a *sub*-nanopore to molecular volume using block co-polymers and A$\beta_{1\text{-}42}$. (a-c)** Selected single blockade currents, $\Delta I$, acquired three block co-polymers *block*-*p*(R)$_{10}$-*co*-(G$_3$S)$_3$-G 23 AAs long (denoted by R-G-S), *block*-*p*(K)$_{10}$-*co*-(G$_3$S)$_3$-G 23 AAs long (denoted by K-G-S) and *block*-*p*(K)$_{24}$-*co*-(A$_8$P$_8$)$_2$-A$_8$ 48 AAs long (denoted by K-A-P), respectively, using the same *sub*-nanopore, which was 0.6 nm in diameter at the waist (ER153). These block co-polymers were used as controls to gauge the read accuracy (from the amplitude) and resolution (from the slope between the blocks) of the *sub*-nanopore using a 250 kS/s sampling rate and a < 100 kHz bandwidth. The empirical data is plotted versus normalized blockade duration for comparison. Juxtaposed with the empirical data is a naïve volume model (assuming $k$ = 1, red line) plotted versus putative acid position assuming a uniform AA velocity through the pore. The data was acquired with a bias voltage of 0.30 – 0.40 V applied across the membrane in electrolyte containing 100 pM of the peptide, 250 mM *NaCl* and 265.5 nM SDS and 1 mM β-ME. The abrupt steps in data shown are commensurate with large volume changes between Arg(R, 0.1734 $nm^3$) and Gly(G, 0.0601 $nm^3$) $\Delta V = 0.1133\, nm^3$, Lys(K, 0.1686 $nm^3$) and Gly $\Delta V = 0.1085\, nm^3$, and Lys(K, 0.1686 $nm^3$) and Ala(A, 0.0878 $nm^3$) $\Delta V = 0.0808\, nm^3$. Likewise, the sharp slope between R and G, and K and G were also consistent with single residue resolution ($k$ = 1). Beyond the steps, the subtle modulations of fluctuation amplitudes between Ser(S, 0.089 $nm^3$) and Gly indicate that differences as small as ($\Delta V = 0.00289\, nm^3$) could be detected this way. However, not all of the blockades showed precisely the same fluctuation patterns. **(d)** The noise power spectral density (*PSD*) acquired when A$\beta_{1\text{-}42}$ was impelled 0.6 V across an *a-Si* membrane nominally 5 nm thick through a *sub*-nanopore (LT076) with a diameter of 0.4 nm defined by the shot noise at the waist. The open pore current was $I_0$ = 220 pA. Superimposed on the plot is a guide (dotted red line) indicating 1/*f* noise. **(e)** A compendium of plots is shown of the average open pore rms-current noise ($\sigma$) measured over a duration of 100 μs within 200 samples of a blockade event for five different *sub*-nanopores. The irreproducible noise diminishes according to a $\sigma \sim 1/\sqrt{N}$ (dotted black line), where $N$ denotes the number of events. **(e, inset)** The average open pore rms-current noise acquired from the LT076 *sub*-nanopore is shown versus time with the number of 150 μs events, $N$, as a parameter. **(f)** A plot is shown of blockade current consensus versus normalized duration measured when denatured SA$\beta_{1\text{-}42}$ was electrically forced through a *sub*-nanopore (PM17, red line). The signals forming the consensus were acquired at 0.6 V amplified over a 1.8

MHz bandwidth, sampled at 3.6 MS/sec and digitally filtered to reveal the fluctuations. Comprising 500 blockades, the consensus signal swings over about 5$\sigma$. Juxtaposed with the empirical data is the corresponding volume model for the peptide (assuming $k$ = 1, blue line). The blockade current was well correlated ($PCC$ = 0.49) to the naïve volume model.

**SUPPLEMENTAL FIGURE 4.**

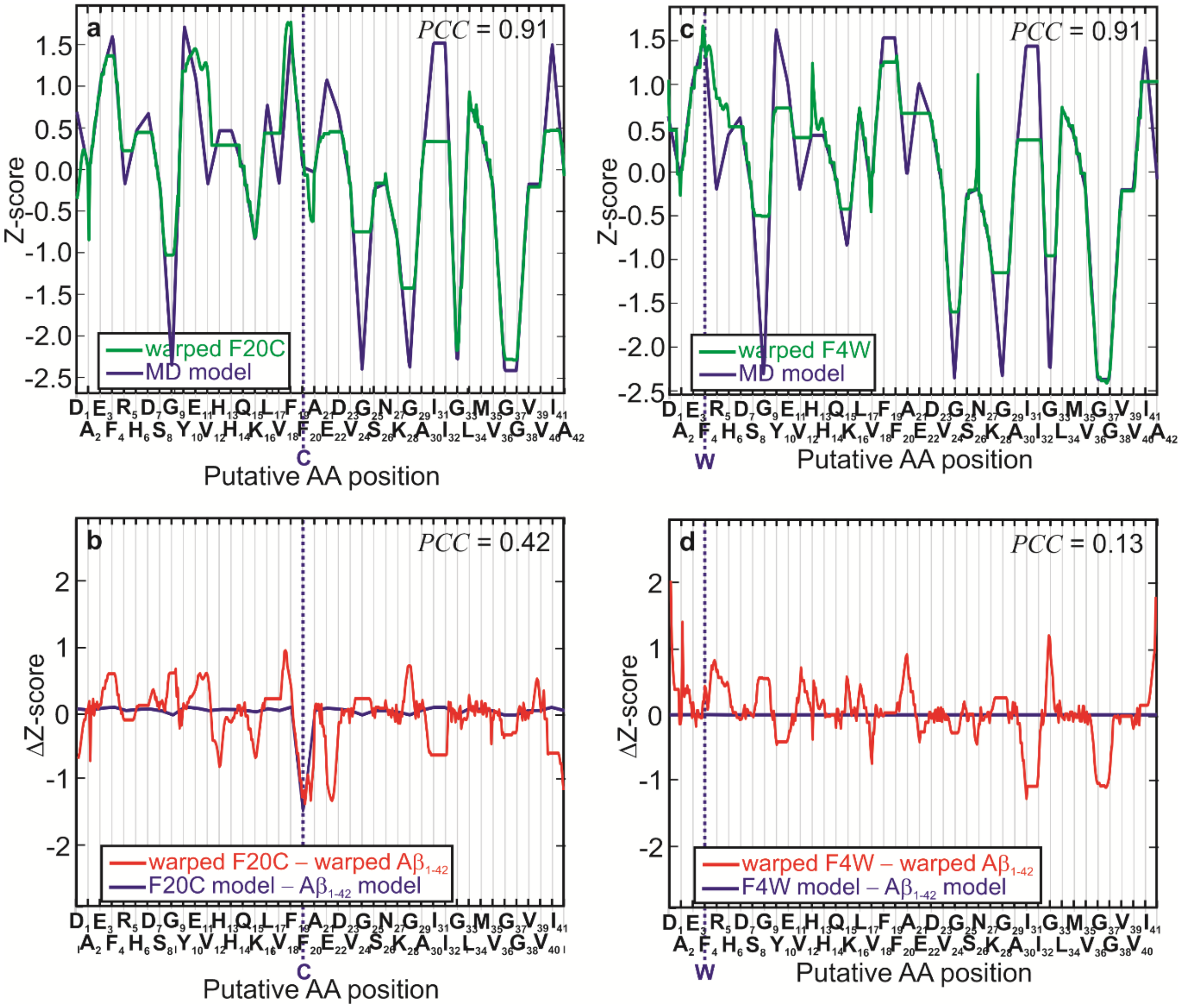


**SUPPLEMENTAL FIGURE 4. Proteoform sequence analysis with near single acid resolution using a *sub*-nanopore. (a,c)** Empirical consensuses associated with blockades due to two mutants of A$\beta_{1\text{-}42}$, F20C and F4W, respectively, are shown after alignment to the corresponding MD models versus the putative AA positions (green lines). Juxtaposed with the empirical consensuses are the corresponding MD models for each variant (blue lines). The dotted blue lines indicate the site of the mutation. The aligned consensuses were all well correlated to the respective volume models with *PCC* ≥ 0.91. **(b,d)** The differences in the Z-scores between the aligned consensus for A$\beta_{1\text{-}42}$ and the respective aligned consensuses for the variants in (a,c) (red lines) are shown versus the putative AA positions. (The A$\beta_{1-42}$ reference data used for comparison—see Fig. 3e—was nearly perfectly aligned with the respective MD model *PCC* = 0.95.) Juxtaposed on the same plots are the differences in the Z-scores between the respective MD models for A$\beta_{1\text{-}42}$ and the corresponding variants (blue lines). The difference between the empirical consensuses were correlated to the difference in the corresponding models (0.13 < *PCC* < 0.42).

**SUPPLEMENTAL FIGURE 5.**

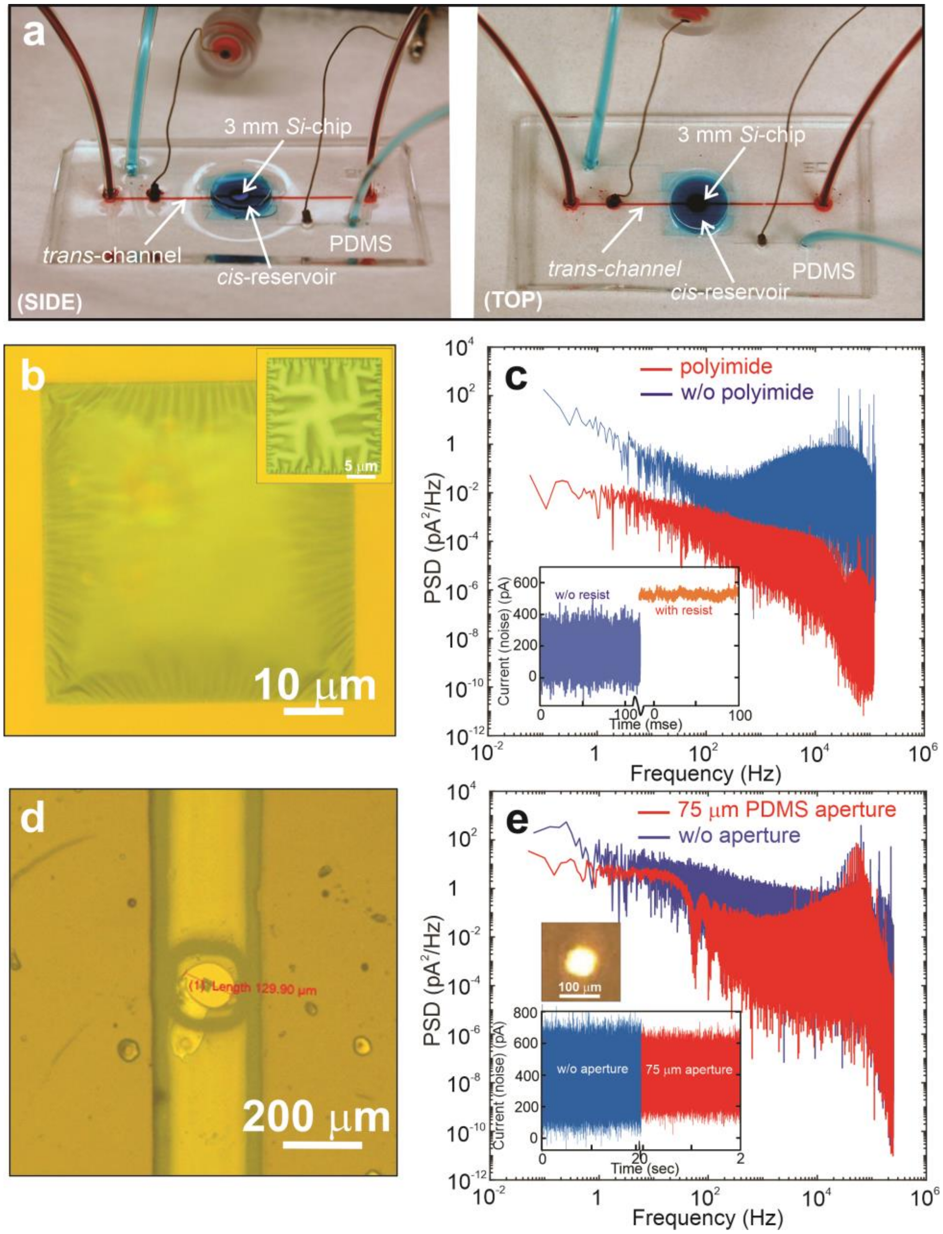

**SUPPLEMENTAL FIGURE 5. Noise mitigation with a *sub*-nanopore through an *a-Si* membrane laminated with polyimide or polydimethylsiloxane (PDMS). (a)** TOP and SIDE views are shown of a microfluidic device supporting a *Si*-chip with a pore through it (Adapted from reference 44). **(b)** An optical micrograph is shown of a 5 nm thick a-*Si* membrane spanning a 50 μm window through a silicon handle wafer with a thick (5 μm) polyimide layer laminated to it to reduce the stray capacitance. **(b, inset).** Like (b), but a smaller area *a-Si* membrane. The polyimide was exposed to UV light through the membrane so the size of the via is the size of the membrane. **(c)** A comparison of the power spectral density (*PSD*) of nanopores through membranes are juxtaposed: one is laminated with polyimide (light blue) and the other is without polyimide (dark blue). **(c, inset)** The current noise is illustrated for *sub*-nanopores with and without a polyimide laminate. The rms current noise with a polyimide laminate was 21.4 pA, whereas without the laminate the noise was 79.7 pA. The dielectric noise (dark blue) along with the amplifier noise predominate at high frequency. (d) An optical micrograph of a silicon chip with a membrane through it bonded to a PDMS microfluidic at the top of a 75 μm via. **(e)** A comparison of the power *PSD* of *sub*-nanopores through 5 nm thick *a-Si* membranes are juxtaposed: one laminated with PDMS (red), another without (dark blue). **(e, inset)** The current noise is illustrated for *sub*-nanopores with and without a PDMS laminate.

**SUPPLEMENTAL FIGURE 6.**

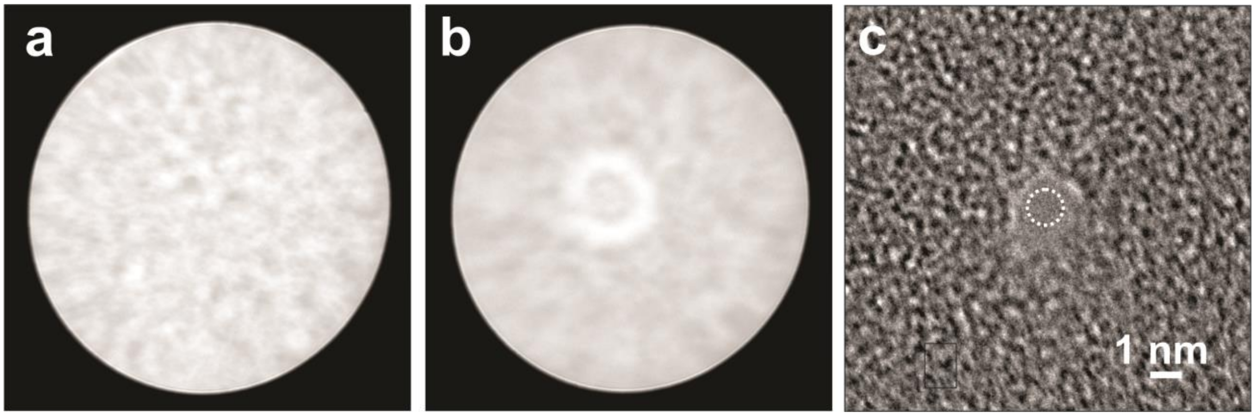


**SUPPLEMENTAL FIGURE 6. Sputtering a pore with a tightly focused, high energy electron beam in a Scanning Transmission Electron Microscope (STEM). (a)** The Ronchigram shown is the diffraction pattern of a convergent beam focused on an amorphous silicon (*a-Si*) membrane specimen that is nominally 5 nm thick. It was obtained with the FEI Titan 80-300 STEM using a beam energy of 300 keV with a spot-size of 6 and a beam current of 466 pA. **(b)** The same Ronchigram in (a) is shown but after exposing the membrane to the beam for 25 sec. **(c)** A TEM image is shown *in vacuo* of a pore immediately after sputtering through the *a-Si* membrane. The cross-section of the pristine pore was estimated from the shot noise associated with electron transmission through the pore to be about 1 nm, but after exposure to the ambient, oxidation likely reduces the lumen to sub-nanometer dimensions.

**SUPPLEMENTAL VIDEOAB1-42E. A molecular dynamics simulation visualizing the translocation of beta-amyloid through a sub-nanopore.**

Following the scheme outlined in the METHODS, visual molecular dynamics (VMD) snapshots are compiled into a video depicting $A\beta_{1\text{-}42}$ translocating across a 5 nm thick silica membrane through a *sub*-nanopore with a bi-conical topography with a cone angle of 8° near the waist, expanding to 36° near each orifice that mimicks the STEM image shown in Fig. 1d. To construct the *sub*-nanopore, initially a crystalline silica slab was generated with a number of unit cells required to fill the pore excluded volume of the amorphous silica ($SiO_2$) membrane with a density of 3.0 g/cm$^3$. The surface geometry of a double conical *sub*-nanopore having an elliptical cross-section was modeled using the relation:

$$\sqrt{\tan^2(\alpha)\cdot x^2+\tan^2(\beta)\cdot y^2} < \sqrt{a\cos^2(\Phi)+b\sin^2(\Phi)\cdot\tan(\alpha)\cdot\tan(\beta)}+|z|\tan(\alpha)\cdot\tan(\beta)+\delta$$

where $\alpha$, $\beta$ are the cone angles at semi-major ($a$) and -minor ($b$) axis of the ellipse (at the minimum cross-section of the *sub*-nanopore near the waist), $\Phi$ is the angle between $x$-axis and a position vector of atoms inside the *sub*-nanopore in $x$-$y$ plane and $\delta$ is the van der Waals radii of the pore surface atoms (taken as 0.16 nm). The *Si* and *O* atoms were randomly placed inside the membrane, excluding the volume of *sub*-nanopore, and then the entire membrane was subjected to simulated annealing a high temperature of 7000 K. A repulsive grid potential was applied to avoid the entry of $SiO_2$ atoms inside the *sub*-nanopore and CHARMM force field parameters were used for *Si* and *O* atoms in the membrane. Later, the bonds between the *Si* and *O* atoms of silica were defined using a 0.22 nm distance cut-off. Using this protocol, a silica *sub*-nanopore having $0.5 \times 0.8$ nm$^2$ minimum cross section was created with a realistic topography (Fig. 1d).

The coordinates of $A\beta_{1\text{-}42}$ (pdbcode:1IYT) protein structure were downloaded from the protein data bank (PDB). The protein structure was placed at the orifice of the pore to study their translocations through the silica *sub*-nanopore and CHARMM36 force field parameters were employed. The constructed silica *sub*-nanopore-protein complex was solvated with a 5 nm thick TIP3P water film on either side of the pore to screen the electrostatic interactions of the pore with its periodic images. The $Na^+$ and $Cl^-$ counter ions were added to the solvated *sub*-nanopore to set 200 mM *NaCl* concentration. Initial systems were subjected to 2000 steps of conjugate-gradient minimization to remove the bad contacts and followed by 1 ns NPT equilibration at temperature 300 K with a 1 fs time step for integration to achieve the bulk water density to 1.0 g/cm$^3$. Periodic boundary conditions were used in all three directions.

A Nośe–Hoover thermostat with a damping coefficient of 1 ps was connected to silica atoms of the membrane to maintain the temperature. Subsequently, the equilibrated systems were subjected to a 20 ns NVT MD simulation with a 0.12 nm cutoff for calculating the short-range part of the non-bonding interactions and the long-range electrostatic interactions were calculated using Particle-Mesh-Ewald (PME) method. The equilibrated *sub*-nanopore system was subjected to an external applied electric field for 25 ns to obtained steady-state ionic current and the last 5 ns trajectory was used to obtained current-voltage characteristics of the *sub*-nanopore.

To simulate the translocation of a peptide, the molecular force fields describing water, ions, and the peptide were combined with the CHARMM force fields for the $SiO_2$ in a 5 nm thick membrane. To mimic the experiments, the membrane separated an aqueous solution of *NaCl* into two compartments connected by a bi-conical *sub*-nanopore with a 0.5×0.8 nm$^2$ cross-section at the waist to match the topography and (linear) current-voltage characteristics (0.194 nS in MD compared to 0.3 nS empirically) of the pore.

At first, electrophoresis was used to force the peptide into the pore alone, but the competition with ions near the membrane and/or non-specific binding of the acids to the surface pinned the molecule in a spot for an extended duration, blocking the ion flow. Alternatively, the peptide was impelled through the *sub*-nanopore by applying a voltage of 25 V across the membrane with a zero-acceleration velocity of $8\times10^{-5}$ nm/ps.
